# Prey diversity revealed by environmental DNA elucidates ecological specialisation and social structure in bottlenose dolphins

**DOI:** 10.64898/2026.06.06.730606

**Authors:** Manuela R. Bizzozzero, Katharina J. Peters, Svenja M. Marfurt, Stephanie L. King, Erik P. Willems, Riccardo Cicciarella, Felix M. Smith, Simon J. Allen, Richard. C. Connor, Michael Krützen

## Abstract

Human disturbance drives rapid biodiversity loss, yet how prey diversity shapes individual niche differentiation and scales up to structure social organisation remains poorly understood, particularly in marine systems where prey are largely invisible to observation. Here we mapped prey community structure (‘prey fields’) using environmental DNA metabarcoding, integrating these maps with stable isotope data and >20 years of behavioural observations on bottlenose dolphins. We show that dolphins specialised on specific prey fields to varying degrees, and that none of 471 tracked individuals changed their predominant prey field — even through climate-driven habitat reorganisation. Isotope profiles confirmed dietary differentiation scaling with specialisation strength. Across four independent sex–subpopulation networks, prey field overlap predicted social affinity in non-foraging contexts, revealing that prey community structure organises social ties beyond foraging. These results expose a socio-ecological link between prey diversity, trophic ecology, and social organisation, providing a transferable eDNA framework for predators with elusive prey.

## Introduction

Biodiversity varies across landscapes and seascapes, creating heterogeneous ecological contexts for individuals. Increasing human pressures are now driving rapid biodiversity declines across terrestrial, freshwater, and marine ecosystems (*1*), yet little is known about how spatial variation in biodiversity structures individual niche differentiation and scales up to shape social organisation (*2*). This is particularly consequential in long-lived, socially complex species where individual ecological strategies can persist across decades, spread culturally, and may shape the social fabric of entire populations. Marine mammals exemplify all these traits, yet socio-ecological studies in marine systems remain rare because prey diversity is difficult to observe directly. Here, we address this gap by using published maps based on an environmental DNA (eDNA) metabarcoding study (*3*) to characterise spatial variation in prey communities and test how prey diversity predicts niche specialisation, diet, and social organisation in Indo-Pacific bottlenose dolphins (*Tursiops aduncus*) in Shark Bay, Western Australia.

In the past, researchers inferred individual foraging niches in marine systems from environmental proxies such as depth, substrate type, or seagrass distribution (*4–6*). While readily available, these variables are often one dimensional and approximate potential prey distributions poorly, as areas that are similar according to such proxies can host fundamentally different prey assemblages. Depth, for example, is often used to delineate microhabitats of cetaceans (e.g., Kopps et al., 2014; O’Brien et al., 2020; Zanardo et al., 2018), yet similar depths can host entirely different prey assemblages depending on salinity or substrate structure (*3*). Without quantifying prey directly, it is difficult to distinguish genuine ecological specialisation from apparent space-use patterns shaped by philopatry, demography, or social preferences. The core problem is that prey biodiversity in marine systems is difficult to observe, making the ecological contexts that directly structure predator behaviour and social structure largely inaccessible.

Advances in molecular biodiversity assessment have begun to change this. Environmental DNA metabarcoding can reveal entire species assemblages from trace genetic material with high spatial resolution across marine, freshwater, and terrestrial systems (*9–11*). When combined with high-resolution remote sensing data, eDNA enables prey diversity patterns to be extrapolated across large areas (*12, 13*). This opens the possibility of defining predator habitats in the same currency predators use to make foraging decisions: spatially delineated prey community structure (i.e., prey fields) rather than environmental proxies — and testing longstanding but empirically inaccessible predictions about how prey diversity structures individual niche differentiation and its effects on social organisation.

Within populations, individuals often diverge in their foraging niches, meaning individuals only exploit a subset of the resources that all individuals in a population collectively use (*14*). Individuals can diverge in niches through a combination of different dietary choices (*15*), by employing different foraging techniques (*16*), or by searching different microhabitats for food (*17*). When individual niche differences arise from consistent behavioural phenotypes, individuals exploiting different niches are termed ‘specialists’ (*14*). Prey diversity is an important factor driving such specialisations (*18*). Empirical studies demonstrate that individual specialisations can persist over months to years, indicating that they often represent stable ecological strategies rather than transient behavioural states (*14*). Furthermore, specialisations occur along a continuum, such that populations may consist entirely of specialists or comprise a mixture of more and less specialised individuals (*19, 20*).

Individual niche differentiation, once established, is not only a pattern of resource use (*14, 21*), but structures encounter rates and association patterns when individuals repeatedly exploit the same resources (*22*). In fission–fusion societies, heterogeneous resource distribution alone can generate structured social networks, as individuals converging on shared patches can form stronger associations (*23*). Such patterns can extend beyond foraging contexts, for example, in a fission–fusion population of Indo-Pacific bottlenose dolphins, individuals specialising on trawler-associated feeding formed distinct social clusters that, after trawling ceased, reintegrated with the wider population, illustrating how changes in feeding ecology can reshape social organisation (*24, 25*). In such socially fluid systems, it is also expected that an individual’s social connectedness can depend on its degree of niche specialisation, with less specialised individuals occupying more central roles in the social network because they traverse a broader set of ecological contexts and social opportunities (*26*).

Cetaceans provide an exceptional model for investigating how prey diversity shapes individual specialisation and social structure. They are highly mobile, socially complex, long-lived and behaviourally flexible, with diverse (and often socially transmitted) foraging techniques (*27–29*). These specialisations are often associated with specific ecological contexts and can shape patterns of social associations (*30–32*). To date, however, individual niche specialisations are easiest to detect when prey differences are conspicuous, such as killer whale (*Orcinus orca*) ecotypes specialising on hunting mammals versus fish (*33*), or when individuals use visible tools or structures, such as the aforementioned dolphins foraging in association with trawlers, leaving specialisations arising from spatially distinct prey community composition largely undetected and understudied.

Shark Bay, Western Australia, provides a rare opportunity to fill this gap. The system hosts the longest-running study of wild Indo-Pacific bottlenose dolphins (*35*). Behavioural data from two subpopulations in ecologically comparable sites in the eastern and western gulfs span more than two decades and more than 20,000 boat-based surveys, in which individually identified dolphins were followed briefly and their GPS location and group activity recorded. The dolphins display remarkable behavioural variation, from widespread foraging techniques such as belly-up fish chasing, ‘snacking’ (*36*), and bottom grubbing in seagrass beds (*37*) to highly specialised, culturally transmitted techniques such as sponge tool use, ‘sponging’ (*38, 39*). Socially, Shark Bay dolphins live in a complex open fission–fusion society with adult males and females socially segregated (40). Males form bonds with other unrelated males in a multi-level alliance network (*35, 41*), whereas female associations are more fluid and often influenced by maternal kinship and habitat similarity (*32, 42, 43*). Dolphins are not territorial, with individuals exhibiting overlapping home ranges that range from 30-160 km_2_ (*8, 35, 44*).

Previous studies in this population suggested that habitat-associated niche segregation predicts social organisation in both sexes (*4, 7, 8, 32*), but because habitats were defined using environmental proxies never linked to prey composition, it remained unclear whether these patterns reflected genuine ecological differentiation or artefacts of space use.

Yet, recently Bizzozzero and colleagues (2025) complemented the long-term behavioural dataset by collecting 270 eDNA samples for metabarcoding of ray-finned fish (Actinopterygii) — the main food source of dolphins in Shark Bay. Using machine-learning models, they delineated five distinct prey fields across both study sites in 2021 (*3*).

In combination, these datasets allowed us to ask whether: (1) individuals specialised on different prey fields and, if so, how strongly; (2) such specialisations were stable across years; (3) prey field specialisation was reflected in diet; and, using social network analysis, how strongly; (4) prey field overlap predicted social associations and (5) specialisation strength predicted individual network position. To reconstruct the ecological context for our long-term behavioural dataset, we hindcasted prey field maps across the study period and calculated individual specialisation indices quantifying how consistently each dolphin used a subset of available prey fields. We linked these specialisations to diet using stable isotope data (δ^13^C and δ^15^N) and to social organisation through network analysis of non-foraging association data.

Together, this work provides a transferable eDNA-based framework for revealing ecological structure hidden from traditional observation and a foundation for comparative socio-ecological analyses in a large-brained, culturally complex marine mammal. Understanding which prey fields individual predators depend on is increasingly critical as accelerating biodiversity loss restructures prey communities worldwide. If prey community structure shapes individual strategies and social networks even in one of the most intensively studied wild mammal populations in the world, analogous structure may be widespread — but has simply remained out of reach.

## Results

### Hindcasting the prey fields

We hindcasted prey field configurations for four additional reference years (2002, 2010, 2014, 2016) using published seagrass maps (*45*) and the machine learning algorithm trained in (*3*). Combined with the published 2021 map, this spanned five key time points (Fig. S1) and allowed us to assign each behavioural survey to its temporally closest prey field configuration, defining five reference periods: ref. period 2002: 1998-2006; ref. period 2010: 2007-2012; ref. period 2014: 2013-2015; ref. period 2016: 2016-2018; ref. period 2021: 2019-2024. Prey field configurations were relatively stable across decades (Fig.1), with the most pronounced decline of prey field A (beige) in 2014 in the western gulf; and in prey field B (green) in 2016 in the eastern gulf (Fig. 1B). Both prey fields, A and B, are based mostly on seagrass and declines correspond to documented seagrass loss following the 2011 marine heatwave.

**Fig. 1.**
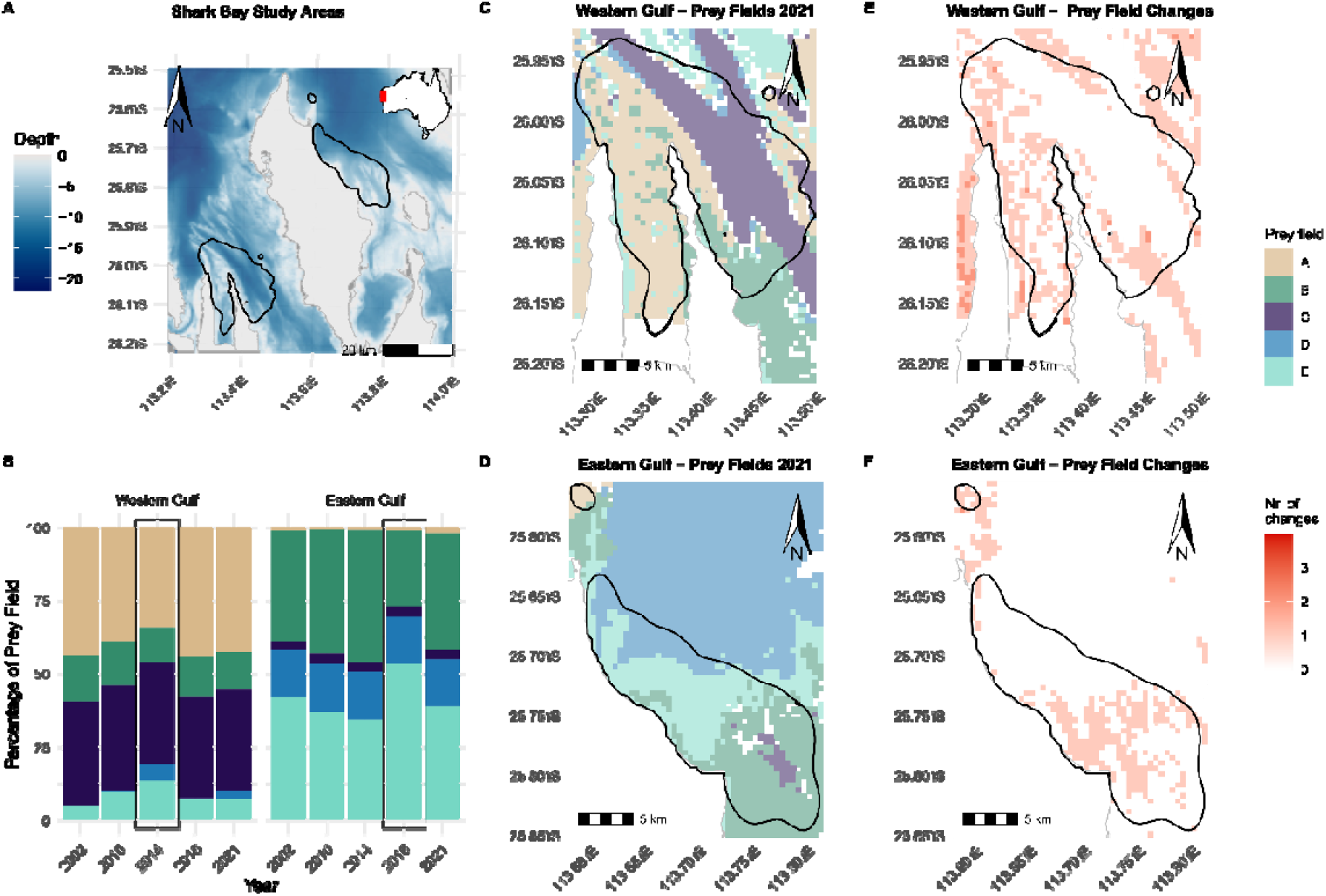
Study sites and spatial distribution of prey fields in Shark Bay, Western Australia. **(A)** Overview of Shark Bay showing bathymetry (depth in metres) and the two Indo-Pacific bottlenose dolphin (*Tursiops aduncus*) study sites (black outlines, 90% kernel density utilisation distributions). Inset shows the location of Shark Bay on the Australian continent (red box). **(B)** Prey field coverage percentage per reference year within each of the two study sites, years with the least seagrass coverage and corresponding prey field configuration change are marked in black rectangles. (**C** and **D**) Spatial distribution of the five prey fields, labelled A–E, in the western and eastern gulf, respectively, as delineated by eDNA metabarcoding of ray-finned fish (Actinopterygii) in 2021 (3). Prey fields were extrapolated across the study sites using a random forest algorithm trained on depth, salinity, seagrass coverage, and channel habitat. Only grid cells with assignment probability ≥ 0.4 are shown in colour, lower values are shown as whitespace. (**E** and **F**) Number of prey field changes across five reference years (2002, 2010, 2014, 2016, 2021) in the western and eastern gulf, respectively. Darker colours indicate cells that shifted prey field identity more frequently across reference years, reflecting seagrass loss following marine heatwave events. Black outlines indicate study site boundaries.

### Dolphins exhibit strong but variable individual specialisation

Using prey field configurations across reference periods, we quantified individual and subpopulation-level specialisation (*S*_*I*_ and *S*_*P*,_ with lower values indicating stronger specialisation) based on a total of 14,151 foraging surveys (western gulf: n_female_ = 1,991; n_male_ = 1,480; n_unknown_ = 122; eastern gulf: *n*_*female*_ = 5,628; *n*_*male*_ = 4,903; *n*_*unknown*_ = 27). We were able to calculate the *S*_*I*_ of 500 individuals, comprising 160 from the western gulf (*N*_*female*_ = 86; *N*_*male*_ = 66; *N*_*unknown*_ = 8; Fig. S2), and 340 from the eastern (*N*_*female*_ = 167; *N*_*male*_ = 171; *N*_*unknown*_ = 2; Fig. S2).

In each subpopulation, low *S*_*P*_ values show that dolphins used on average only a fraction of the locally available and collectively used prey fields, indicating that each subpopulation consisted predominantly of specialised individuals. This pattern was statistically significant compared to a null model accounting for sampling effort and data structure: western gulf *S*_*p*_ = 0.48, p < 0.001; eastern gulf *S*_*p*_ = 0.69, *p* < 0.001 (Fig. S3).

Consistent with the expected continuum of specialisation, individuals varied markedly in their degree of specialisation (Fig. 2). Individual *S*_*I*_ values ranged from 0 to 1.13 and from 0 to 1.36 in the western and eastern gulfs, respectively. An individual specialisation index of zero indicates an individual foraged exclusively in one prey field; higher values indicate progressively more even prey-field use, with *S*_*I*_ equal to one corresponding to a within-individual diversity (Shannon entropy) equal to that of the pooled subpopulation. Values above 1 arise when a dolphin distributes its foraging more evenly across prey fields than the subpopulation does collectively. Individual foraging proportions across prey fields at the base of these indices are shown in Fig S4.

**Fig. 2.**
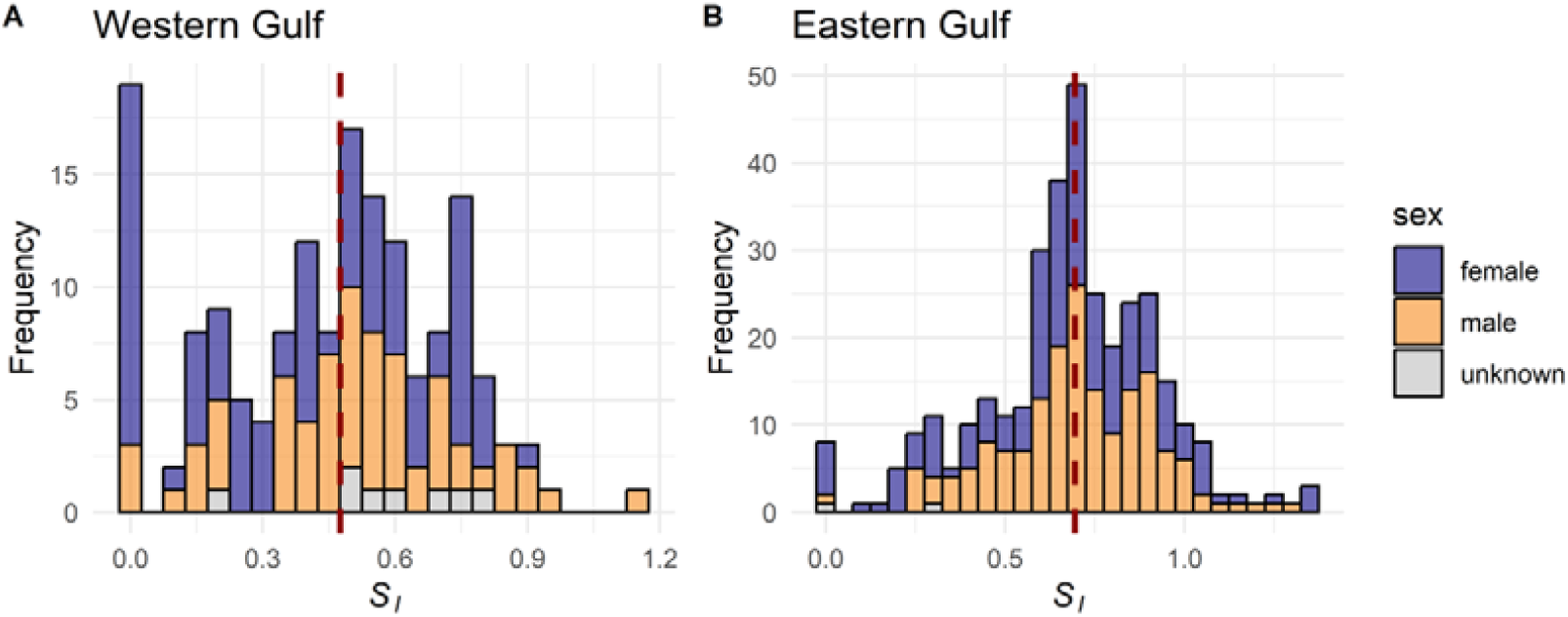
Distribution of individual foraging specialisation indices (*S*_*I*_). Bars are colour-coded according to sex. **(A)** Shows individuals of the western and **(B)** of the eastern gulf subpopulations. Subpopulation-level specialisation indices (*S*_*p*_) are indicated with a red dashed line.

Of the 160 observed individuals in the western gulf, 58.1% predominately foraged in prey field A; 37.5%, 3.8%, 0.6% and 0% in prey fields C, B, E and D, respectively. This loosely traced average prey field availability over all reference periods in this study site (A: 40.6%, C:35.2%, B:13.9%, E: 8.3% and D: 2.0%; Fig. 1B).

Of the 340 observed individuals in the eastern gulf, 333 could be assigned a single predominant prey field; the remaining seven showed ties between two fields. Of those 333, 54.7% predominately foraged in prey field E; 42.6%, 2.1%, 0.6%, and 0% in prey fields B, C, D, and A, respectively. This loosely traced average prey field availability across all reference periods in this study site (E: 41.1%, B: 38.3%, D: 16.2%, C: 3.3%, and A: 1.1%; Fig. 1B). Prey field D was the exception: it covered 16.2% of the eastern gulf on average, yet only 2 of 333 individuals specialised on it.

### Specialisation strength varies with sex and subpopulation

The specialisation strength differed systematically across subpopulation and sex (Fig. 3). A Bayesian generalised linear model (GLM) incorporating these variables explained 18% of the variance in *S*_*I*_ (*R*^*2*^ _*c*_ = 0.180, 95% CI = 0.13–0.23) and substantially outperformed the null model including only sampling effort (expected log predictive density difference *ΔELPD ± SE* = −42.3 ± 9.0). After excluding individuals of unknown sex, the model included data on 167 and 86 females and 171 and 66 males from the eastern and western gulf subpopulations, respectively.

**Fig. 3.**
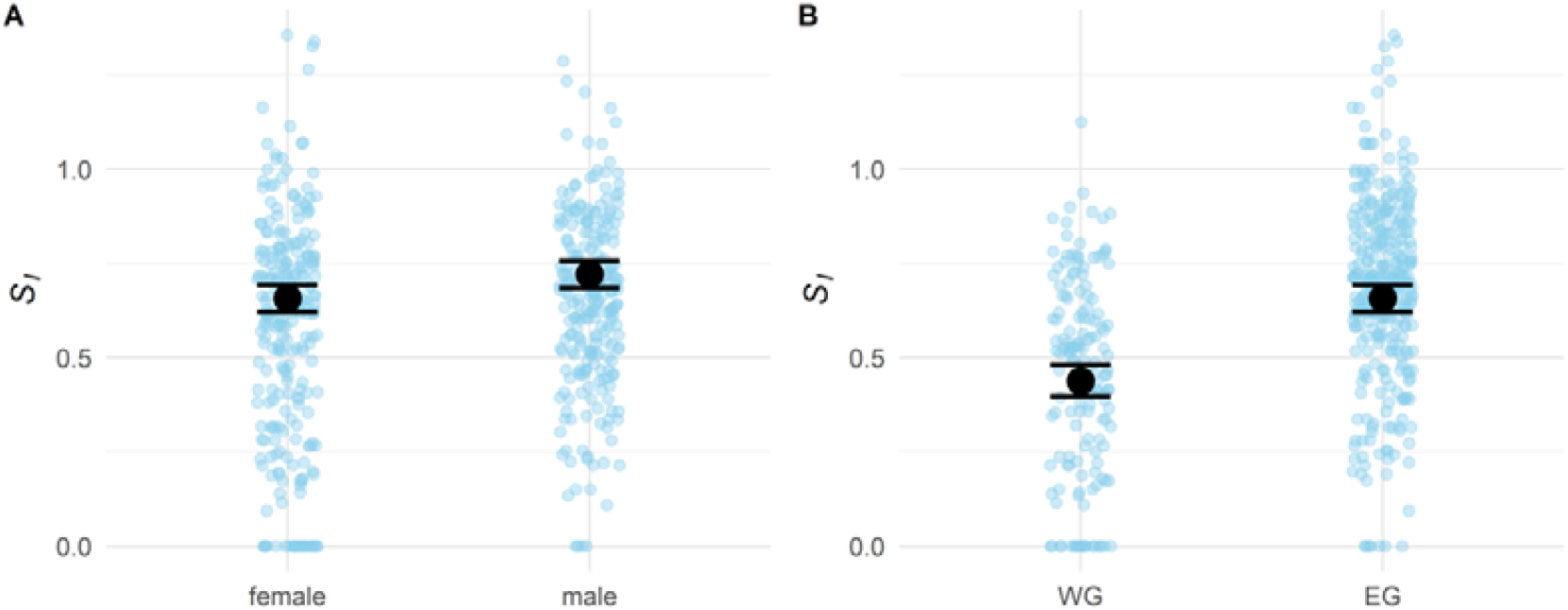
Conditional effect plots of estimated effects of sex and subpopulation on *S*_*I*_ values. Light blue points represent individual observed data points, while black points and whiskers indicate posterior means and their 95% credible intervals. **(A)** displays the relationship between *S*_*I*_ and sex, **(B)** illustrates the effect of (WG: western gulf subpopulation, EG: eastern gulf subpopulation).

Sex differences were modest. Males showed slightly higher *S*_*I*_ values (*β* = 0.07, 95% CI = 0.02 – 0.11) than females, indicating marginally lower specialisation (Fig. 3A). Individuals from the subpopulation in the western gulf had consistently lower *S*_*I*_ values, i.e., were more specialised, than those in the eastern gulf (*β = −0*.*25*, 95% *CI = −0*.*31 – −0*.*19;* Fig. 3B). Variation in *S*_*I*_ attributable to differences in sampling effort was small but accounted for (log(surveys): *β = 0*.*05*, 95*% CI = 0*.*01* – 0.09). Including individual age had no effect on *S*_*I*_ (SI Section S1, Table S1).

### Foraging specialisations persist through climate-driven habitat change

To evaluate the temporal consistency of individual specialisation, we determined each individual’s predominant foraging prey field for each reference period. After restricting analyses to individuals observed in at least two reference periods, we examined 471 dolphins for temporal consistency in predominant prey field (western gulf: *N*_*females*_ = 81; *N*_*males*_ = 65; *N*_*unknown*_ = 8; eastern gulf: *N*_*females*_ = 154; *N*_*males*_ = 161; *N*_*unknown*_ = 2;).

None of the 471 individuals included in this analysis changed their predominant prey field across reference periods, indicating striking long-term stability in prey field specialisation (Fig. 4). This stability persisted despite spanning multiple decades of observation and extensive prey field configuration shifts (Fig. 1E & F), especially in 2014 (western gulf) and 2016 (eastern gulf).

**Fig. 4.**
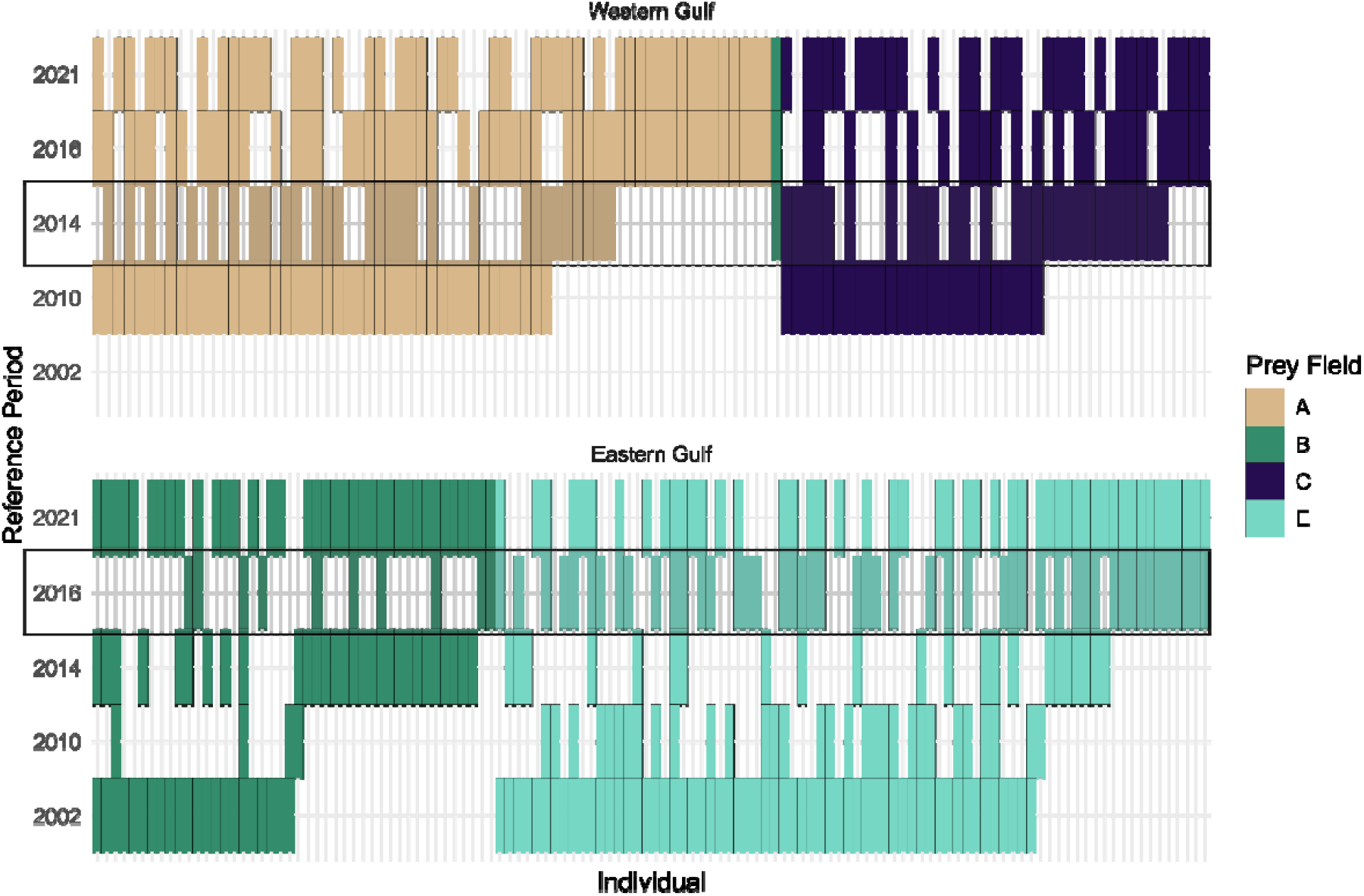
Temporal stability of individual foraging habitat preferences. Per subpopulation, each row represents a reference period, while each column corresponds to an individual. Coloured tiles indicate the predominant foraging prey field for each individual in a given period. Reference periods with most drastic habitat reconfiguration due to seagrass coverage changes are highlighted with black rectangles.

### Prey field specialisations are reflected in trophic differentiation

To assess whether prey field specialisation was reflected in trophic differentiation, we analysed previously published stable isotope data from dolphin biopsies (*34, 46*). Nitrogen isotope ratios (δ^15^N) indicate trophic position, whereas carbon isotope ratios (δ^13^C) reflect the primary production sources supporting the food web. In total, we analysed 14 tissue samples from the western gulf and 21 from the eastern gulf subpopulations (Fig. S5).

After confirming no interannual trend in either isotope (SI Section S2, Table S2), we tested whether δ^15^N differed among individuals specialising in different prey fields. A Bayesian GLM model including predominant prey field and the potentially confounding variables sex and subpopulation substantially outperformed the null intercept-only model (*ΔELPD ± SE* = −9.1 ± 3.9; *R*^*2*^_*c*_ = 0.51, 95% *CI* 0.31 – 0.64). Sex showed a modest effect (*β* = 0.23, 95% *CI* 0.03 – 0.43), while subpopulation (*β* = −0.26, 95% *CI* −5.89 – 5.48) had non-credible to marginal effects.

Since individual specialisation strength itself covaries with sex and subpopulation, including all predictors simultaneously would introduce redundancy. We therefore tested a complementary model where δ^15^N differentiation among prey fields was modulated by individual specialisation strength, fitting a model with the interaction between predominant prey field and *S*_*I*_. This interaction captures the expectation that isotopic distinctiveness reflects both prey field identity and the strength to which individuals restrict their foraging to that field. The interaction model credibly outperformed the null (*ΔELPD ± SE* = −12.9 ± 4.9) and marginally outperformed the sex–subpopulation model (*ΔELPD ± SE* = −3.8 ± 2.9), explaining 65% of variance in δ^15^N (*R*^*2*^_*c*_ = 0.65, 95% *CI* 0.49 – 0.74). We retained the interaction model as the primary analysis because it slightly outperformed the sex–subpopulation model in predictive accuracy. This suggests that the interaction between predominant prey field and *S*_*I*_ captures the variation associated with sex and subpopulation, while also adding biologically meaningful information beyond it.

Relative to prey field A, individuals specialising in prey field C showed markedly elevated δ^15^N (*β* = 2.05, 95% *CI* 1.33 – 2.77), and those in prey field B moderately elevated values (*β* = 1.13, 95% *CI* 0.43 – 1.85). Both prey fields showed strong negative interactions with *S*_*I*_ (prey field B × *S*_*I*_: *β* = −1.71, 95% *CI* −3.00 – −0.44; prey field C × *S*_*I*_: *β* = −2.49, 95% *CI* −3.92 – −1.05), indicating that more specialised individuals exhibited increasingly distinct trophic signatures within their prey field (Fig. 5). Prey field E estimates were highly uncertain (*β* = −0.15, 95% *CI* −2.23 – 1.79; prey field E × *S*_*I*_: *β* = −0.07, 95% *CI* −2.59 – 2.68), reflecting the small sample size of individuals predominantly foraging in this prey field (*N* = 5). δ^13^C values showed no detectable effect of prey field or specialisation (SI Section S3, Table S3, Fig. S6), suggesting prey fields do not differ substantially in basal carbon sources.

**Fig. 5.**
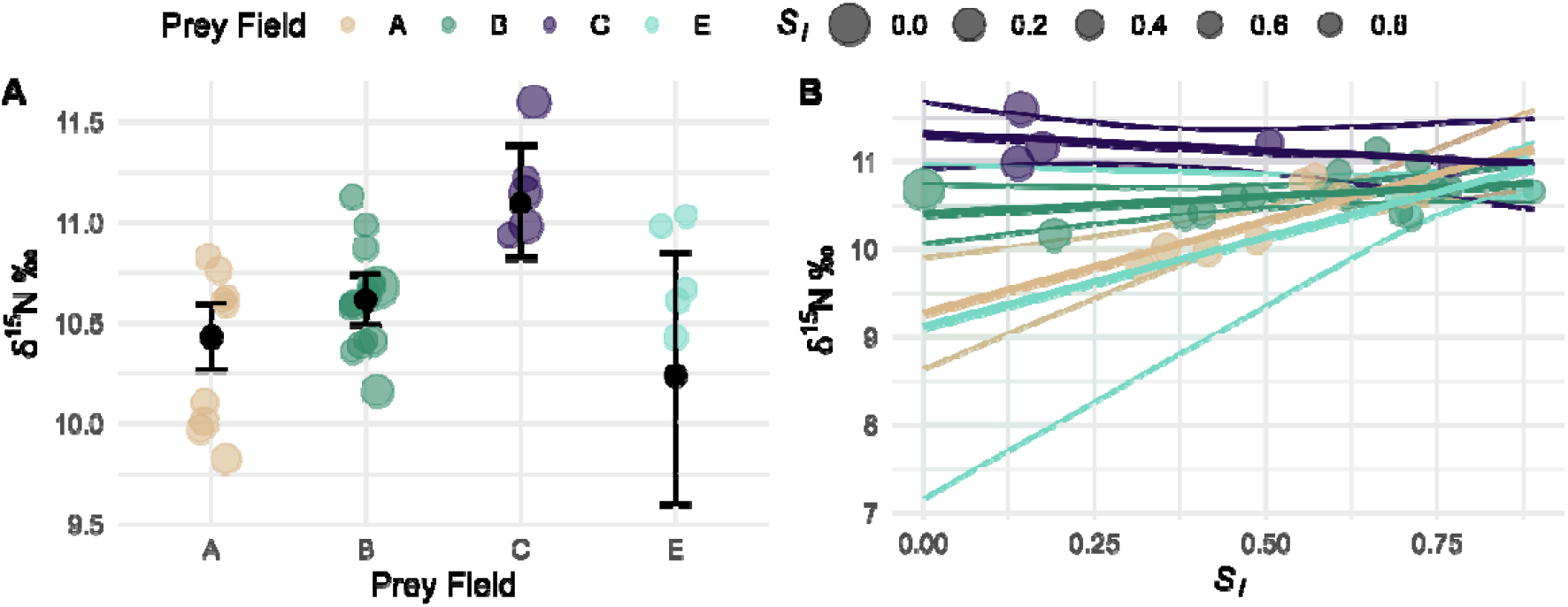
Relationship between prey field specialisation and δ^15^N ratios. Coloured points represent observed δ^15^N ratios of individuals and their predominant prey field. The size of the point indicates the strength of specialisation S_I._ Both panels show predictions from the same model with predominant foraging prey field and *S*_*I*_ as predictors. **(A)** Black points with 95% credible interval error bars show the conditional effect of prey field, averaged across *S*_*I*_ values. **(B)** Coloured lines with 95% credible intervals show the interactive effect of prey field and *S*_*I*_, illustrating how the relationship between *S*_*I*_ and δ^15^N differs between prey fields.

### Individuals specialising on the same prey fields cluster socially outside foraging contexts

Individuals with greater overlap in foraging prey fields spent more time together outside of foraging contexts. Across all four sex–subpopulation combinations, pairwise prey field overlap (*PS*_I_) significantly predicted social affinity, quantified by the Simple Ratio Index (*SRI*). All model comparisons strongly supported the niche specialisation model over the null model accounting only for individual-level variation in association rates (Table 1, Fig. 6, Fig. S7). The strength of this relationship was greatest among western gulf females, followed by eastern gulf females, and then western and eastern gulf males. Hierarchical clustering of social association matrices further revealed prey field identity as an underlying organising principle of social structure, with individuals sharing prey fields forming deeply rooted social clusters — also most pronounced in the western gulf and among females (Fig. S8).

**Table 1.**
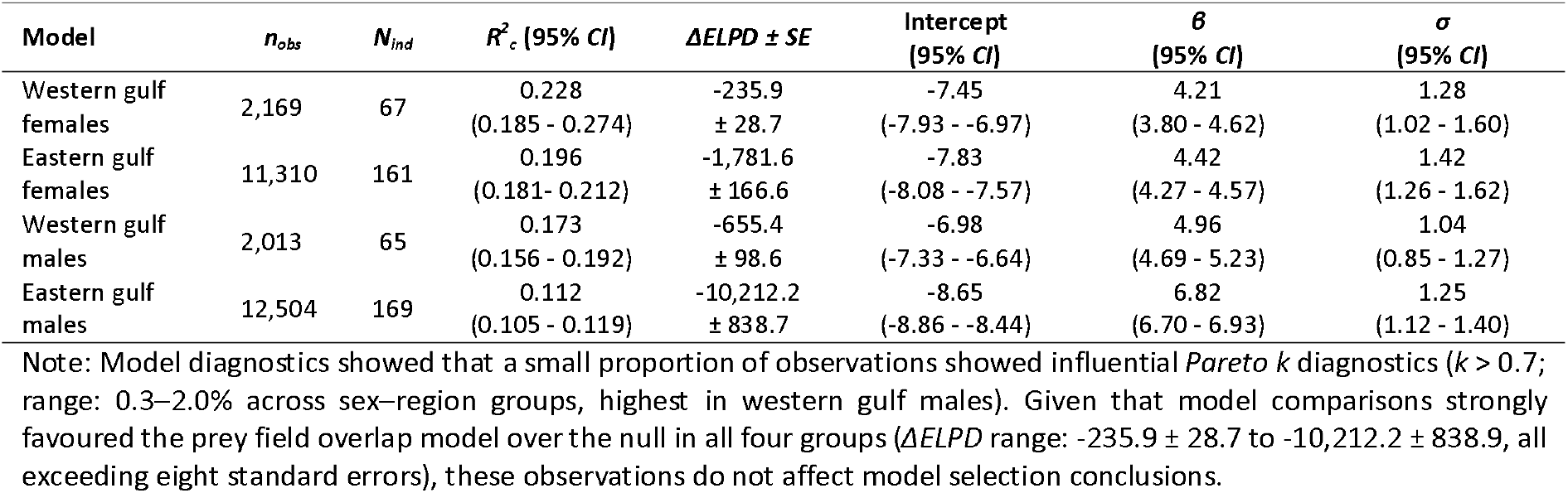
Summary of the results of the social network analyses. Summary table of Bayesian zero-inflated binomial GLMM computed for the four separate datasets consisting of eastern and western gulf females and males, respectively. *n*_*obs*_ describes the number of dyads included in the analyses and *N*_*ind*_ the number of individuals included. ΔELPD ± SE describes the difference between the null model only including the random intercept for each individual within dyads, thereby accounting for individual-level variation in association rates, and the model including the *PS*_*I*_ of a dyad as a predictor. Subpopulations and sexes are ordered according to variances explained by the model in decreasing order.

**Fig. 6.**
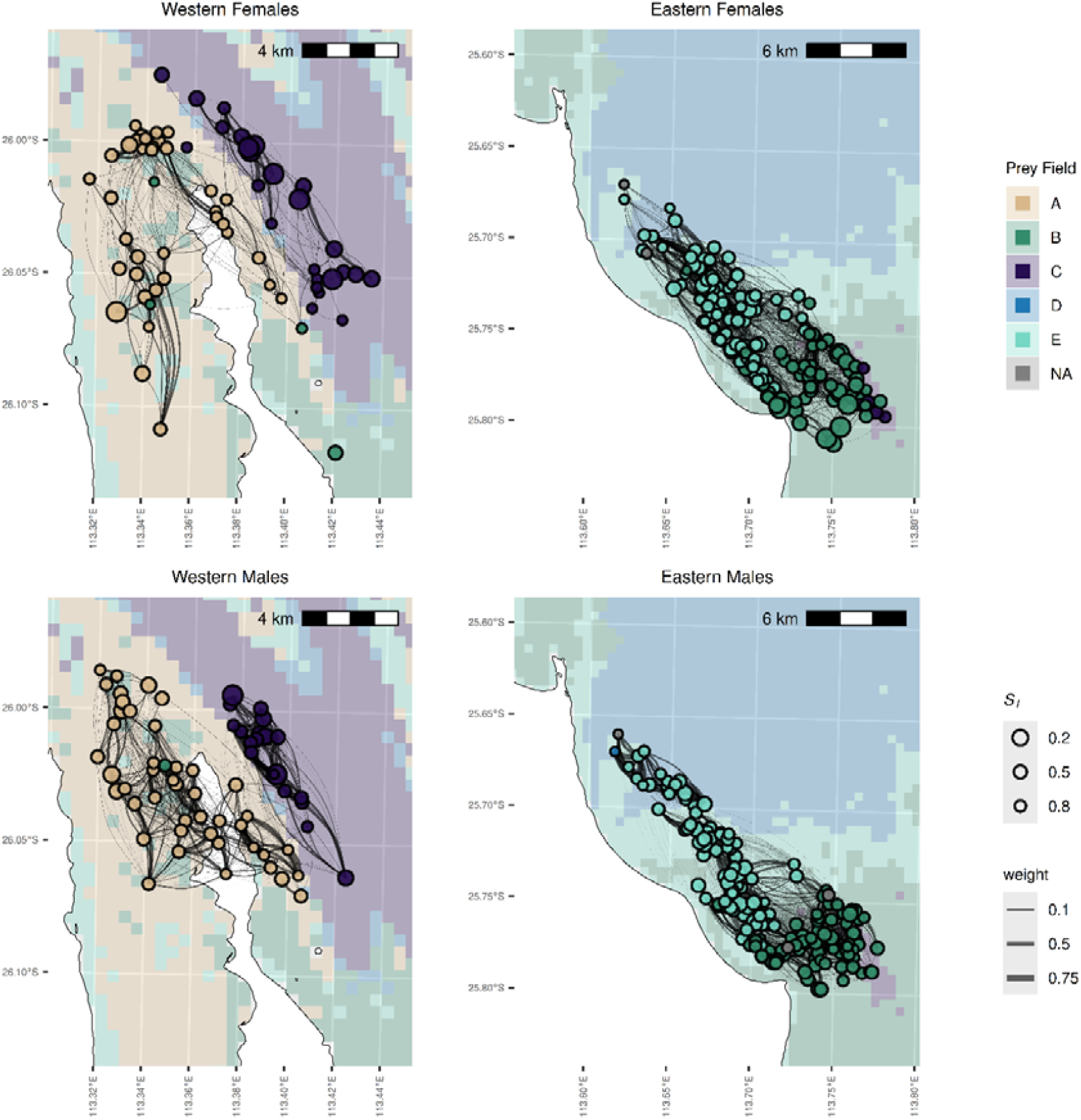
Social network plots for females and males from the eastern and western gulfs of Shark Bay. Individual Indo-Pacific bottlenose dolphins (*Tursiops aduncus*) are represented as network nodes (*points*), which are connected by edges (*lines*) indicating social associations. Nodes are coloured according to the individual’s predominant foraging prey field and are sized based on the individual’s strength of specialisation *S*_*I*_. Their positions correspond to the centroid of their foraging surveys. Edge widths represent dyadic social relationship indices *SRIs*, reflecting the strength of associations between individuals in non-foraging surveys. The background map displays prey fields using coloured tiles according to the prey field map of the 2021 reference period, while landmasses are shown in white.

### Sex shapes social network position more than foraging specialisation

We quantified each dolphin’s social position using four centrality metrics derived from non-foraging association networks built separately for each sex and subpopulation (463 individuals across four networks: western gulf females = 67, western gulf males = 65, eastern gulf females = 161, eastern gulf males = 169): degree (number of associates), strength (summed association indices), mean association strength (average bond strength per partner), and betweenness (frequency of bridging otherwise disconnected individuals). We predicted that foraging generalists would show broader, less committed social ties — higher degree and strength, lower mean association strength, and higher betweenness.

Across all three joint models, the most consistent pattern was a sex difference in how individuals are socially positioned within their networks (Table 2). Mean association strength was credibly higher in males than females, and males also tended toward lower total association weight (i.e., strength), while showing no credible difference in number of associates (i.e., degree). Read together, this is consistent with males and females occupying structurally different social systems: males maintaining stronger average bonds across a comparable number of partners, females distributing comparable association activity across more partners with weaker average bonds per partner.

**Table 2.**
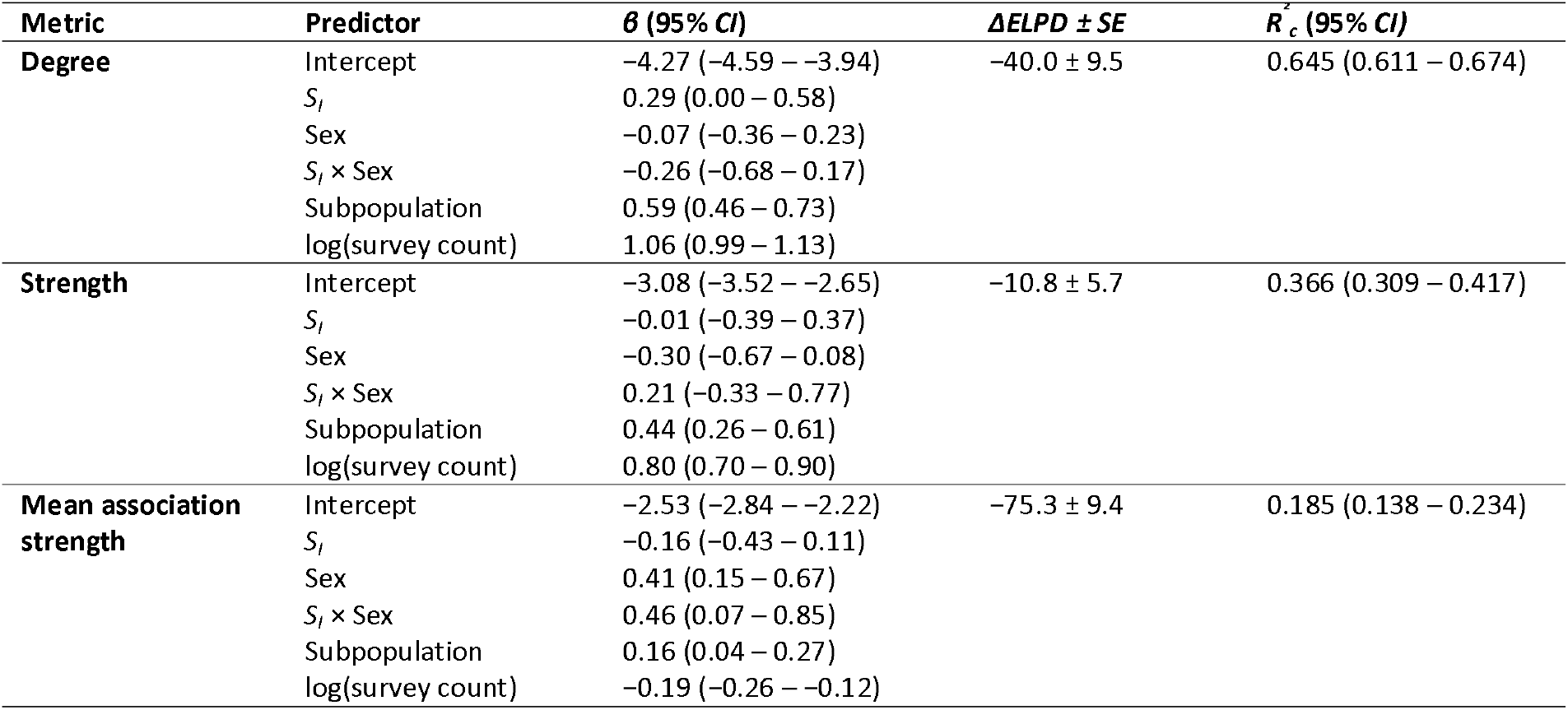
Summary of the Bayesian models linking foraging specialisation to centrality in non-foraging social networks. Degree and strength were standardised within network prior to modelling and analysed with Gaussian families; mean association strength was analysed on the raw scale with a Beta family (logit link). All three models were fitted jointly across both subpopulations with *S*_*I*_, sex, their interaction, subpopulation, and log-transformed non-foraging sighting frequency as predictors. Each model was compared via leave-one-out cross-validation against a null model including only log-transformed sighting frequency; *ΔELPD ± SE* and Bayesian *R*^*2*^ _*c*_ (with 95% *CI*) refer to the focal model. *β* values are posterior means with 95% credible intervals; the reference level for sex is female and for subpopulation is eastern gulf. Metrics are ordered by *R*^*2*^ _*c*_ in decreasing order.

Within this sex-structured backdrop, foraging specialisation appeared as a weaker secondary modifier of social position. For degree centrality, more generalist individuals tended to have more recorded associates, however, with the lower credible bound reaching zero, this is best interpreted as a suggestive trend rather than a credibly non-zero effect. The S_I_ main effect on strength was effectively null, so the trend in degree did not translate into proportionally greater total association weight. The most credible effect of specialisation was its interaction with sex on mean association strength, indicating that more specialised males had stronger average bonds per partner, whereas, in females, the relationship was flat to weakly negative. Across all four sex–subpopulation networks, betweenness centrality models did not outperform their respective null models, and specialisation showed no credible association with betweenness in either the gamma or hurdle component of any subgroup (Table S4).

## Discussion

By defining prey fields from measured fish community structure rather than environmental proxies, we could test socio-ecological predictions that have remained difficult to evaluate empirically in marine systems. Doing so revealed that Indo-Pacific bottlenose dolphins consistently specialise on distinct prey fields, that these specialisations persist over decades — including through a period of substantial habitat reorganisation — and that ecological similarity predicts social affinity outside of foraging contexts. Documented across two demographically independent subpopulations and over 20 years of behavioural data, these results suggest that spatial prey community structure is reflected in individual niche differentiation and social organisation.

### Prey-defined seascapes reveal strong and persistent individual specialisation

Using Roughgarden’s framework (47), we showed that individuals exploited a smaller subset of the realised prey fields than the collective use of prey fields of each subpopulation in each gulf, reinforcing the idea that individuals within a population are not ecologically equivalent (*14*). The specialisation strength occurred along a continuum, ranging from highly specialised individuals foraging exclusively in one prey field to less specialised foragers that nonetheless exhibited clear preferences. This pattern matches continua documented in other systems, such as pigeon guillemots, *Cepphus columba* (*48*), and bluegill sunfish, *Lepomis macrochirus* (*49*). In marine mammals, the documentation of such continua are limited to a few studies using stable isotope analyses, e.g., in Australian fur seals, *Arctocephalus pusillus doriferus* (*50*), or in species with visible prey items, e.g., sea otters, *Enhydra lutris nereis* (*51*). Our framework now allows the investigation of individual specialisation in a system where prey is invisible to direct observation in a non-invasive manner.

Importantly, individual specialisations were remarkably stable over time. Across both gulfs, dolphins consistently specialised in the same prey field over multiple reference periods. This specialisation persistence occurred even after dramatic habitat changes driven by a significant marine heatwave event in 2011 triggering seagrass die-offs which impacted dolphin survival and reproduction (*39*) and regrowth during the subsequent years (*45*), and a configuration shift from prey field A to E in the western gulf in 2014 and B to E in the eastern gulf in 2016. Rather than shifting into newly formed prey fields within the same spatial area, specialists of prey field B were observed less frequently within our core survey zone following its reconfiguration, suggesting that they tracked their preferred prey assemblages beyond the study boundaries.

This pattern is consistent with distinct ecological preferences rather than with passive site fidelity alone. These findings align with the ‘distinct preference model’ of (*52*), in which individuals develop first-choice resources (here prey fields) and maintain them due to strong behavioural trade-offs. In long-lived mammals with socially transmitted techniques, such trade-offs may be reinforced by learning constraints and energetic investment (*29, 53*). Frequency-dependent trade-offs may further stabilise specialisation (*54*). Persistent specialisation therefore likely arises from the interaction of learning constraints, behavioural trade-offs, and frequency-dependent dynamics.

Stable and persistent specialisations mirror long-term specialisation observed in bottlenose dolphins at population levels at larger geographic scales (*32, 55*). The observation that habitat-associated differentiation operates consistently across spatial scales, from ocean basins to shared foraging grounds within a single population, suggests that ecological fidelity to prey communities may be a fundamental and scale-independent organising principle in this species, with implications for how population-level divergence originates and is maintained.

### Individual specialisation is reflected in trophic differentiation

Stable isotope analyses revealed that prey fields captured ecologically meaningful trophic differences. Individuals specialising in different prey fields showed distinct δ^15^N signatures, with trophic divergence scaling with specialisation strength, but similar δ^13^C ratios across prey fields, consistent with a shared nearshore carbon baseline (*56*). Highly specialised individuals showed the most pronounced isotopic differentiation, whereas generalists exhibited more intermediate isotopic profiles. These findings demonstrate that our eDNA based approach captured the realised ecological niches of individual dolphins rather than abstract abiotic spatial categories. Traditional trophic tools such as stable isotopes, fatty acids, or stomach contents, reveal dietary composition (*57*), but cannot delineate the spatial organisation of prey assemblages (*58*). By contrast, prey-defined seascapes provide an explicit ecological map linking resource structure to individual behaviour.

In systems where prey species are cryptic, mobile, or difficult to sample, our approach provides a powerful alternative for linking behaviour to trophic ecology. More broadly, it suggests that individual-level niche variation can be quantified without assuming fixed prey identities, using prey community structure as an integrative ecological measure.

### From ecological similarity to social structure

Ecological similarity predicted social affinity: by quantifying prey fields directly rather than inferring habitat from proxies, we show that ecological structure predicts substantial components of social organisation in a highly mobile marine mammal. Dolphins foraging in the same prey field associated more frequently outside of foraging contexts. This is despite them living in an open, unbounded fission–fusion society where group composition changes frequently and individuals have ample opportunity to associate broadly (*35*). Our results are consistent with the idea that repeated convergence on shared ecological contexts can structure encounter rates and, over time, social networks (*59, 60*). Our study thus provides empirical support for resource dispersion theory (*22*) and its extensions to dynamic social systems (*23*). Such patterns were suspected based on studies using environmental approximations for prey distributions (*5, 6, 32*), but, until now, were rarely explicitly linked to ecological preferences.

Remarkably, social structuring was not binary but graded. Social segregation reflecting niche differentiation was strongest in the subpopulation and sex with higher degrees of overall specialisations, i.e., in the western gulf and in females. This suggests that the strength of specialisation could shape the strength of social segregation. Since specialisation strength depends on biodiversity patterns (*61, 62*), these findings elucidate a mechanical pathway through which biodiversity patterns can shape social organisations.

At the level of individual network position, sex was a more consistent organising axis than specialisation, with males maintaining credibly stronger average bonds per partner than females while showing comparable numbers of associates. This pattern is broadly consistent with the well-documented male alliance system (*63– 65*). Within this sex-structured backdrop, specialisation acted as a weaker secondary modifier: more generalist individuals showed a suggestive trend toward more recorded associates, and the relationship between specialisation and average bond strength differed credibly between sexes, with more specialised males maintaining stronger average bonds; a pattern absent in females. This represents a rare empirical test on wild animals of theoretical predictions that ecological specialisation can structure social connectivity in group-living species (*26, 66*). Together, these subpopulation- and individual-level patterns suggest that the continuum of specialisation, rather than its presence alone, shapes the social fabric of animal societies.

### Ecological structuring and social feedbacks: the limitations of our approach

Our data cannot disentangle whether ecological differentiation preceded social clustering, or inversely, whether pre-existing social groups converged on shared foraging strategies. Critically, however, both mechanisms implicate ecological context as a necessary condition. If ecological similarity drives social affinity, prey field overlap directly structures who associates with whom. If pre-existing social groups converge on shared strategies, the convergence itself requires an ecological substrate: shared prey fields must exist for groups to specialise on them collectively. Social structure and ecological context are therefore not independent regardless of causal direction, and prey community structure remains a fundamental organising axis of dolphin societies under either scenario.

The male social system illustrates this point particularly clearly. Male bottlenose dolphins in Shark Bay form multi-level alliance networks structured around cooperative mate access, with adult partner choice predicted by affiliation history and age similarity rather than kinship (*35, 41, 65*). Shared foraging tactics have long been suspected to contribute to alliance formation, as coordination is more tractable among males with similar foraging behaviour (*8, 30*). Our results provide direct evidence for this: even within a social system organised primarily around mating cooperation, males preferentially associated with others sharing similar foraging specialisations. This suggests that ecological preferences constitute a persistent and fundamental axis of social structuring that operates alongside, and partially independently of other social forces: not overriding alliance dynamics, but detectable even within a system where those dynamics dominate.

### Bottlenose dolphins as a model for eco-social and evolutionary processes

Bottlenose dolphins offer a uniquely powerful model system for investigating how ecological heterogeneity structures social systems. Unlike systems characterised by near-complete social segregation (e.g., killer whale ecotypes), dolphin societies exhibit partial, flexible segregation: specialists preferentially associate with one another but remain socially connected. This intermediate structure mirrors early-stage divergence processes, where ecological specialisation generates social structuring without complete reproductive isolation. Such systems provide fertile ground for studying cultural niche construction and gene–culture co-evolution (*7, 67*).

### A general framework for predators with elusive prey: Implications for conservation

Beyond dolphins, this study provides a generalisable roadmap for integrating eDNA-derived prey fields with long-term behavioural data. Where long-term behavioural records are unavailable, the same framework can be applied to telemetry or other movement data. Many predators forage on prey that are difficult or impossible to observe directly, including pelagic fishes, deep-sea mammals, nocturnal terrestrial carnivores, and aerial insectivores. In such systems, prey fields derived from eDNA and environmental extrapolation offer a scalable, non-invasive means of quantifying ecological niches at biologically relevant scales; and, as our results illustrate, of revealing individual-level structure that is invisible when habitat is approximated by abiotic proxies alone.

This matters for conservation because individual specialisation can promote population-level resilience by maintaining behavioural and ecological diversity (*26, 68*), yet the same specialisations may also increase vulnerability when specialised resources decline and specialists cannot flexibly shift (*14, 69–71*). Our data show that such specialisations can be deeply entrenched: over two decades and through documented climate-driven seagrass loss, no single tracked individual switched its predominant prey field. Understanding where, how, and by whom resources are exploited is therefore critical for effective conservation planning (*72*). For apex predators, preserving not only population size but also ecological diversity may be key to maintaining ecosystem stability and preventing trophic cascades (*73*).

Prey-informed maps are particularly well suited to this task. They identify which areas support distinct prey assemblages, can be monitored over time, and can be modelled under future climate-change scenarios. They can also reveal which prey communities support — or fail to support — individual specialisation in the first place. For example, the almost complete absence of specialists in open-water prey field D, despite its substantial area, suggests that predictable, benthic or seagrass-associated prey fields B, C, and E offer more favourable conditions for specialisation than schooling fish in pelagic waters (*3, 14, 69*). This kind of asymmetry, between habitat that is available and habitat that actually structures individual ecology, is invisible to area-based assessments alone, but has direct implications for conservation planning and tourist management in systems where managers need to know which habitats sustain ecological diversity rather than simply which habitats are present.

In conclusion, defining predator habitats in the same currency animals use to make foraging decisions — prey community structure rather than environmental proxies — makes long-standing socio-ecological questions tractable at a resolution rarely achieved in marine systems. Across two demographically independent subpopulations and more than two decades of observation, Indo-Pacific bottlenose dolphins consistently specialised on distinct prey fields, maintained those specialisations through climate-driven habitat reorganisation, and socially aligned with ecologically similar conspecifics outside of foraging contexts. That such a fundamental axis of individual variation was invisible until prey were mapped directly underscores how much ecological structure can remain hidden in systems where prey are inaccessible to observation. As prey distributions shift under ongoing environmental change, prey-informed approaches will be increasingly critical for anticipating how biodiversity loss translates into altered ecological strategies, social dynamics, and ultimately population resilience — not only in dolphins, but across the many predator systems where the ecological substrate of behaviour has, until now, been largely out of reach.

## Methods

### Experimental Design

This study was an observational, integrative analysis testing how prey community structure relates to individual specialisation, diet, and social organisation in Indo-Pacific bottlenose dolphins, Tursiops aduncus (*74*). It combined three data streams: environmental DNA-derived prey-field maps generated in a previous study (*3*), more than two decades of behavioural surveys collected in two demographically mostly independent subpopulations in the western and eastern gulfs of Shark Bay, Western Australia, and published stable isotope data from skin biopsies. The units of investigation were individually identified dolphins followed across multiple years.

The study addressed five pre-specified questions, set out at the end of the Introduction. No interventions were applied; all data derive from non-invasive observation and pre-existing biopsy archives. Detailed analyses, together with their inclusion criteria, spatial frameworks, and statistical models, are described in the subsections below.

### Study System

Our two long-term study sites for dolphin behaviour (Shark Bay Dolphin Research) are in the eastern gulf (~320 km^*2*^) and the western gulf (~230 km^*2*^) of Shark Bay, Western Australia (Fig. 1). We defined our core study areas using kernel density estimation at the 90% utilisation distribution level, based on all surveys (see Long-term behavioural data) conducted between 1998 and 2024 in eastern gulf and between 2007 and 2024 in western gulf. Specifically, we used the kernelUD of the ‘adehabitatHR’ package (*75*) function, with the “href” bandwidth selection method (*h* = “href”) and the Epanechnikov kernel (*kern* = “epa”).

### Long-term behavioural data

We used long-term behavioural data collected mostly during the cooler months between May and November, including 15,000 behavioural surveys in the eastern gulf and 6,093 in the western gulf (Fig. S9). During a boat-based survey, we identified dolphins by photographing their dorsal fins (76). Within the first five minutes of a survey, we recorded group size and composition, GPS position, sea state, and predominant group activity (rest, forage, travel, socialise, or unknown) following (*37*) (see ethogram in (*30*)). Group membership was defined using the 10 m chain rule, in which individuals within 10 m of any other group member are considered part of the same group (*37*). We determined an individual’s sex through behavioural observations, i.e., presence of a dependent calf in the infant position (*77*), genital area inspection, or genetic analysis (see also laboratory protocols in (*30*). Biopsy samples were collected opportunistically using a Paxarms biopsy rifle (*78*).

### Prey fields from previous eDNA study

We defined prey fields based on a study that characterised fish distribution across both dolphin study sites in western gulf and eastern gulf during survey efforts in the austral winter of 2021 (3). In brief, via eDNA metabarcoding, the study analysed 270 water samples collected from 45 locations, targeting ray-finned fish (Actinopterygii) specific 16S and 12S rRNA gene regions. The study identified five distinct fish communities (A–E), whose distributions were influenced by four environmental variables: depth, salinity, seagrass coverage, and channel habitat, i.e., deep areas (15-30 m) with complex rocky substrate delineated by steep slopes (*79*). The study used a random forest algorithm to extrapolate these fish communities across the study sites on a 500 × 500 m raster grid, providing a high-resolution framework for defining prey fields. Unlike proxy-based approaches that define habitats directly from environmental variables without explicitly linking them to prey distribution, this approach defines prey fields from directly measured prey community structure and uses explicitly linked environmental correlates solely to extrapolate these prey fields in space (*3*).

Prey fields A and B were located in shallow seagrass habitats, distinguished by varying seagrass coverage, with prey field A occurring in areas with lower seagrass coverage, lower salinity, but higher fish richness than prey field B. Prey field C, located in deeper channel areas with no seagrass, exhibited the highest fish richness. Prey field D occurred in deep and open waters with sandy sea floor and consisted primarily of pelagic schooling fish but had the lowest fish richness. Prey field E occurred in medium-depth area with no seagrass but predominant turf substrate, and with intermediate fish richness (3).

### Hindcasting prey fields

One of the key environmental layers influencing prey field distribution was seagrass coverage. However, the extent of seagrass has been significantly affected by an unprecedented marine heatwave in recent years (*45*), therefore causing potential shifts in prey field composition. To incorporate historical prey field changes into our long-term behavioural dataset, we employed the trained random forest algorithm previously used for contemporary prey field modelling (*3*) to hindcast prey fields for years with available data on seagrass variation. Specifically, documented seagrass extents in Shark Bay for 2002, 2010, 2014, and 2016 (*45*), enabling us to model prey fields under these distinct conditions. For each predicted layer, we evaluated assignment probabilities to prey fields, and only retained assignments with a probability ≥ 0.4 for analysis. Higher thresholds disproportionately excluded cells in the southern eastern gulf, where prediction certainty was lower, causing surveys conducted in these areas to be discarded for downstream analyses (see also sensitivity analysis in Fig. S10). As individuals foraging across multiple prey fields, including these ambiguous zones, would lose surveys selectively, stricter thresholds would artificially inflate specialisation indices by retaining only observations from high-certainty areas. A threshold of 0.4 was therefore chosen as a conservative compromise that maximised spatial coverage while maintaining reasonable assignment confidence. Repeating analyses with a more noise inducing threshold of 0.3 did not significantly impact the results.

These hindcasted prey fields enabled us to assess long-term ecological changes and provided a temporal framework for behavioural analyses. We assigned behavioural data to the prey field of the temporally closest reference year; if two reference years were equidistant, we selected the earlier one. This approach categorised the data into reference periods based on reference years: Ref. period 2002: 1998-2006; Ref. period 2010: 2007-2012; Ref. period 2014: 2013-2015; Ref. period 2016: 2016-2018; Ref. period 2021: 2019-2024.

### Calculation of prey field specialisation indices (S)

We quantified individual prey field specialisation using Roughgarden’s framework (*47*), which partitions the total niche width (*TNW*) of a population into within-individual (*WIC*) and between-individual components. Essentially, this quantification compares the resource use of an individual (*WIC*) to the resource use of every individual in a population collectively (*TNW*). We defined the degree of specialisation, following (*80*), as the proportion of TNW explained by *WIC*:

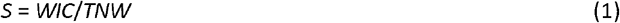

where smaller *S* values denote stronger specialisation. We estimated *WIC* and *TNW* variances using the Shannon-Weaver diversity index (*81*), incorporating the proportion of each individual’s foraging surveys in each prey field (*WIC*), and the proportion of the subpopulations’ collective surveys in each prey field (*TNW*). We calculated S on two levels: the individual level *S*_*I*_, yielding an index for every dolphin; and the subpopulation level *S*_*p*_, yielding an index for each subpopulation in each gulf. The subpopulation level index parametrised the weighted mean degree of individual prey field specialisation in each gulf. More details on these calculations are provided in SI Section S4.

A central assumption of the Roughgarden’s framework in eq. (1) is that all individuals in a population have access to the full set of resources used collectively by the population, i.e., that the population’s realised niche represents the available niche from which individuals select. Several features of our study system support this assumption. First, the two study sites are spatially compact: at the documented cost-of-transport cruising speed of ~2 m s^−1^ for bottlenose dolphins (*82*), an individual can traverse the long axis of either study area in approximately 2–3 hours. Second, dolphins in Shark Bay are not territorial: home ranges of 30–160 km^*2*^ overlap broadly across individuals, with no defended boundaries (*8, 35, 44*). Third, there are no physical barriers (coastal topography, depth, or hydrographic features) that would systematically exclude any individual from any prey field within either study site. Fourth, we have occasionally observed individuals crossing from the western gulf to the eastern gulf study site, thus travelling ~120 km between sites, far exceeding the within-site distances that any individual would need to cover to access all prey fields in its study site. We therefore treat each subpopulation’s collective prey-field use as a reasonable representation of the available prey-field set for each individual, with home-range area itself representing the spatial expression of an individual’s prey-field selection rather than a constraint upon it (*83*).

### Stable isotope samples

For our analyses, we leveraged previously published stable isotope data (δ^13^C and δ^15^N) measured from skin tissue biopsies of dolphins collected at our study sites between 1997 and 2009 (34, 46). In the field, samples were stored frozen in liquid nitrogen, or in saturated NaCl and 20% dimethyl-sulfoxide (DMSO) solution at −20°C. In the laboratory, samples were stored at −80°C until they were lipid-extracted, dried, ground into powder, and analysed as specified in (*46*).

Since the samples were collected over a span of thirteen years, we applied a correction of –0.022‰ year^-1^ (*84*) to the carbon isotope values of dolphin samples. This accounted for changes in δ^13^C values of atmospheric CO_2_ caused by fossil fuel combustion, known as the ‘Suess effect’ (*84, 85*). Given that samples included here ranged from 1997 to 2009, we corrected all carbon isotopic values of samples referenced to the year 2009.

### Calculation of dyadic prey field overlap (PS_I_) and dyadic associations (SRI)

To investigate whether individuals foraging in the same prey field form social subgroups during non-foraging activities, we quantified dyadic prey field overlap and social affiliations between dolphins using established indices: as a measure of pairwise prey field overlap, we used the Proportional Similarity Index (*PS*_*I*_) (*80*), while social affiliations were assessed using the Simple Ratio Index (*SRI*) (*86, 87*). All dyadic measures were calculated separately for each sex and subpopulation (four networks in total).

We used a custom R script to calculate the *PS*_*I*_ and the package ‘asnipe’ (*88*) to calculate the SRI. Detailed index calculations are available in SI Sections S5 & S6.

### Calculation and standardisation of social network metrics

To test whether individual foraging specialisation predicts social network position, we computed four centrality metrics from the previously calculated non-foraging association networks: degree centrality (number of recorded associates), strength centrality (sum of association indices across all associates), mean association strength (strength divided by degree, reflecting the average association index per partner), and betweenness centrality (frequency with which an individual lies on shortest paths between other network members). Together, these metrics capture complementary aspects of social position: degree and strength reflect the breadth and intensity of social connectivity; mean association strength captures bond commitment per partner; and betweenness captures the role of individuals as bridges between otherwise weakly connected parts of the network. We predicted that more generalist foragers (higher *S*_*I*_) would occupy more central network positions; specifically, higher degree, higher strength, lower mean association strength (weaker average bonds distributed broadly), and higher betweenness, because their foraging behaviour is not confined to prey-field-specific contexts shared with specific individuals, facilitating broader and less committed social associations.

Social networks metrics were calculated separately for each sex and subpopulation (four networks in total). Centrality metrics were computed using the igraph package in R (*89*). For betweenness centrality, edge weights were inverted prior to calculation (weights = 1/*SRI*) because igraph interprets weights as distances rather than connection strengths, ensuring that stronger associations were treated as shorter paths.

Degree and strength centrality were standardised within each network (z-scored to *mean* = 0, *SD* = 1) prior to modelling, as their absolute values depend on network size and structure and are therefore not directly comparable across networks. Mean association strength (Strength/Degree) was analysed on the raw scale because dividing by degree removes the network-size dependence of the sum-based strength metric, leaving values interpretable as the average co-occurrence rate per partner regardless of network size. Betweenness centrality was used on the raw scale separately for each network, as its distributional shape differed between subpopulations in ways not addressable by standardisation alone.

### Data inclusion criteria

For the calculation of *S*_*I*_ and *S*_*P*_, we included only foraging surveys of weaned, independent individuals, i.e., above the age of four (*90*). To ensure independent foraging bouts, we retained only one foraging survey per individual per day. We assigned each survey to a prey field using its GPS location and the corresponding reference prey field map based on the date of the survey. We only considered surveys with an assigned prey field and included only individuals with at least 10 foraging surveys to ensure sufficient data for meaningful proportion calculations.

To assess whether individual prey field specialisation was consistent over time, we first assigned each individual a predominant foraging prey field for each reference period. This was based on the same data restriction criteria as for calculating *S*_*I*_, except that we allowed multiple surveys per day, excluding so-called ‘resights’- i.e., surveys of the same individual within one hour - to ensure a sufficient number of observations per reference period per individual. Analyses were further limited to individuals with an assigned foraging prey field in at least two reference periods; reference periods in which an individual was observed fewer than five times were excluded.

To assess whether individual prey field specialisations are reflected in trophic differentiation, we only included data from dolphins for which we had: biopsy samples; assigned a predominant prey field; and calculated a specialisation index *S*_*I*_. As no individual changed its predominant prey field across reference periods (see Result 2), prey-field assignments and *S*_*I*_ values were independent of the temporal window used and could be matched directly to isotope samples regardless of biopsy collection date.

For dyadic prey field overlap, we calculated the *PS*_*I*_ based on the same foraging surveys per individual that were previously used to estimate *S*_*I*_. The SRI was calculated based on non-foraging surveys, including only one survey per day to avoid pseudo-replication. We restricted the analysis to independent individuals of known sex that had been recorded in at least 10 non-foraging surveys. Additionally, to ensure comparability between indices, we included only dolphin dyads for which both *PS*_*I*_ and SRI could be computed under the given data restrictions. We excluded dyads where individuals could not have associated due to a lack of temporal overlap.

Social network metrics were based on the same criteria used for SRI calculation with the exception that one western gulf female (REX_2) was excluded prior to network metric calculation as she had zero recorded associations with any other individual in the dataset, most likely reflecting incomplete spatial coverage at the boundary of the study area rather than true social isolation.

### Statistical Analysis

Specialisation indices were tested against null distributions using Monte Carlo permutation (*80*). All other analyses used Bayesian regression models in the ‘brms’ package (*91*), each compared via leave-one-out cross-validation (LOO-CV) against null models containing only relevant nuisance covariates. Weakly regularising priors were used throughout. For all Bayesian models, we required effective sample sizes above 1,000 and 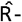 values below 1.01 to confirm adequate posterior sampling, increasing adapt_delta up to 0.99 when necessary. Influential observations were assessed using LOO-CV (*92*); observations with *k* > 0.7 are reported per model where relevant. Model performance was evaluated through graphical posterior predictive checks and quantified using a Bayesian *R*^*2*^_*c*_ statistic (*93*). The detailed statistical models are described in the subsections below.

### Testing for prey field specialisation

To test whether in each subpopulation, on average, dolphins used a significantly smaller portion of the prey field than expected by chance, we implemented a non-parametric Monte Carlo resampling procedure, as proposed by (*80*). In this approach, the observed *S*_*p*_ is compared to a distribution of random *S*_*p*_*s*. To this end, for each gulf, we randomly reassigned prey fields across all surveys conducted within the same reference period and recalculated *S*_*p*_. By preserving the original prey field assignment distribution, each random reassignment reflected the probability of occurrences in the observed dataset. We repeated this process 9,999 times, generating a null model representing a population surveyed with the same effort as the observed data, but with individuals foraging randomly within the population’s realised niche. We then compared the observed *S*_*p*_ values against this null distribution and assessed their significance at a 5% probability level. If *S*_*p*_ exceeded what was expected by chance, we assumed individual specialisation occurred and we assigned each individual to its predominant foraging prey field, defined as the prey field in which it foraged most frequently. All these analyses were performed using a custom R script.

To assess whether individual degrees of specialisation (*S*_*I*_) were significantly affected by sex and subpopulation, we excluded all individuals of unknown sex and implemented a Bayesian Generalised Linear Model (GLM) with the response variable *S*_*I*_ modelled as a truncated Gaussian distribution (trunc(lb=0)) to account for non-negative values. The model included sex and subpopulation; and log-transformed sighting frequency as a nuisance covariate to account for a potential relationship between observation effort and *S*_*I*_. The model was compared against a null model including only log-transformed sighting frequency using LOO-CV.

To aid model convergence, we applied weakly regularising priors, specifying normal distributions for the fixed effects (*μ* = 0, *σ* = 10) and the intercept (*μ* = 0, *σ* = 10). Additionally, we used a Student’s t-distribution (*ν* = 3, *μ* = 0, *σ* = 10) for the residual standard deviation to allow for more robust estimation. We ran the model using MCMC and four chains, each with 2,000 iterations (including 1,000 warm-up iterations).

We repeated this analysis, incorporating an individual’s mean age across all its foraging surveys to test for age effects. However, this approach restricted the analysis to individuals with accurate age estimates, significantly reducing our dataset for the western gulf. Therefore, we presented these results separately in SI Section S1.

### Prey field specialisation stability over years

We examined the number of transitions in predominant foraging prey fields across reference periods to assess the temporal consistency of individual prey field specialisation. However, due to complete lack of variation (i.e., no individuals changed their predominant foraging prey field), statistical modelling of prey field shifts was not appropriate.

We used predominant prey field rather than *S*_*I*_ for temporal comparisons because the number of foraging surveys per individual within a single reference period was often too low to yield reliable proportion-based indices. Predominant prey field assignment is robust to small sample sizes because it depends only on the rank order of prey field use, not on exact proportions.

### Linking prey field specialisation to trophic differentiation

After ruling out interannual trends in isotope ratios (SI Section S2), we fitted a series of Bayesian linear models to δ^15^N and δ^13^C, with Gaussian response distributions, comparing: (i) a null intercept-only model; (ii) a model including predominant prey field, sex, and subpopulation, to assess isotopic differentiation among prey fields while accounting for potential demographic and spatial confounds; and (iii) a model including the interaction between predominant prey field and *S*_*I*_, to test whether isotopic distinctiveness depended on both prey field identity and specialisation strength.

To aid model convergence, we applied weakly regularising priors, specifying normal distributions (*μ* = 0, *σ* = 5) for the fixed effects, normal distributions (*μ* = 0, *σ* = 10) for the intercepts, and Student’s t-distributions (*ν* = 3, *μ* = 0, *σ* = 10) for the residual standard deviations to account for potential outliers. We fitted the model using MCMC sampling with four chains, each run for 2,000 iterations (including 1,000 warm-up iterations).

### Linking prey field specialisation to social structure

To analyse the effect of foraging prey field overlap (*PS*_*I*_) on association strength (*SRI*), we fitted a Bayesian zero-inflated binomial general linear mixed-effect model (GLMM), as per (*32*). To account for the dyadic nature of the data, we specified random effects using a multi-membership structure to control for repeated observations of individuals across dyads. Specifically, we included a random intercept for each individual within dyads, thereby accounting for individual-level variation in association rates. Since females and males have different social strategies and to avoid over-parameterising the model, we ran separate models for each sex and subpopulation. We compared a null model including only the random intercept against a model additionally including *PS*_*I*_ as a predictor.

To aid model convergence in the models for the western gulf subpopulation, we applied weakly regularising priors, specifying normal distributions (*μ* = 0, *σ* = 2) for the intercept, normal distributions (*μ* = 0, *σ* = 1) for the fixed effects, and Cauchy distributions (*μ* = 0, *σ* = 1) for the standard deviations of the random effects. A normal prior (*μ* = 0, *σ* = 2) was placed on the zero-inflation parameter to constrain estimates of excess zero observations. The models were fitted using MCMC sampling with eight chains each run for 8,000 iterations (including 4,000 warm-up iterations).

For the models on the eastern gulf subpopulation, due to low effective sampling sizes, we increased the number of iterations per chain to 15,000 for the male model (using eight chains) and 10,000 for the female model (using six chains). In both models, we regularised the prior of the intercept slightly more (*μ* = 0, *σ* = 1).

### Linking prey field specialisation to social network position

For degree and strength, Bayesian Gaussian models were fitted jointly across both subpopulations, with *S*_*I*_, sex, their interaction, subpopulation, and log-transformed non-foraging sighting frequency (log(IDtotal)) as predictors. The interaction between *S*_*I*_ and sex was included because social organisation differs substantially between sexes in this population. Log-transformed sighting frequency was included as a nuisance covariate to account for the positive relationship between observation effort and recorded association rates. For mean association strength, a Beta family with logit link was used, appropriate for proportional data strictly bounded on the open unit interval. For betweenness centrality, 85 individuals (18.4%) had zero betweenness, indicating they did not act as network bridges. A hurdle Gamma family was therefore used for each of the four networks separately, simultaneously modelling the probability of having non-zero betweenness (hurdle component) and the magnitude of betweenness among bridge individuals (Gamma component). Both components included *S*_*I*_ and log-transformed sighting frequency as predictors. All models were compared against null models including only log-transformed sighting frequency using LOO-CV.

Weakly regularising priors were specified as normal distributions (*μ* = 0, *σ* = 1) for fixed effects, normal distributions for intercepts (*μ* = 0; *σ* = 1 for eastern-gulf networks, *σ* = 2 for western-gulf networks) and exponential distributions (λ = 1) for residual standard deviations in Gaussian models, gamma distributions (*σ* = 0.01, *β* = 0.01) for the shape and precision parameters of Gamma and Beta families, respectively, and normal distributions (*μ* = 0, *σ* = 1) for hurdle component parameters. All models were fitted using MCMC sampling with four chains, each run for 4,000 iterations, including 2,000 warm-up iterations.

## Acknowledgments

This research was carried out in Gathaagudu, Malgana Country, and we acknowledge the Traditional Owners of the region. We thank RAC Monkey Mia Dolphin Resort, Shark Bay Resources, the Useless Loop community, especially Joel Adele, and Jade Standen, Liam Ridgley and Brendan Lumley of Aristocat 2 charters, as well as the Department of Biodiversity, Conservation and Attractions, Shark Bay Rangers for their continued support and assistance. We thank the many research assistants, students, and volunteers who have contributed to long-term data collection in Shark Bay since 1998, without whom this research would not have been possible. We thank Michael Heithaus for openly sharing stable isotope data from his previous research.

## Use of Artificial Intelligence tools

AI assistance (Claude Sonnet 4.6, Anthropic, PBC, accessed via claude.ai, 2025–2026) was used for refining, correcting, editing, and formatting text to improve clarity and flow, as well as for refining and debugging R scripts. AI assistance was not used to create ideas or generate, analyse, or interpret data or images.

