## Supplementary Material for "Prey diversity revealed by environmental DNA elucidates ecological specialisation and social structure in bottlenose dolphins"

Manuela R. Bizzozzero *et al.*

**This PDF file includes:**

Supplementary Text Section S1 - S6
Figs. S1 to S11
Tables S1 to S4

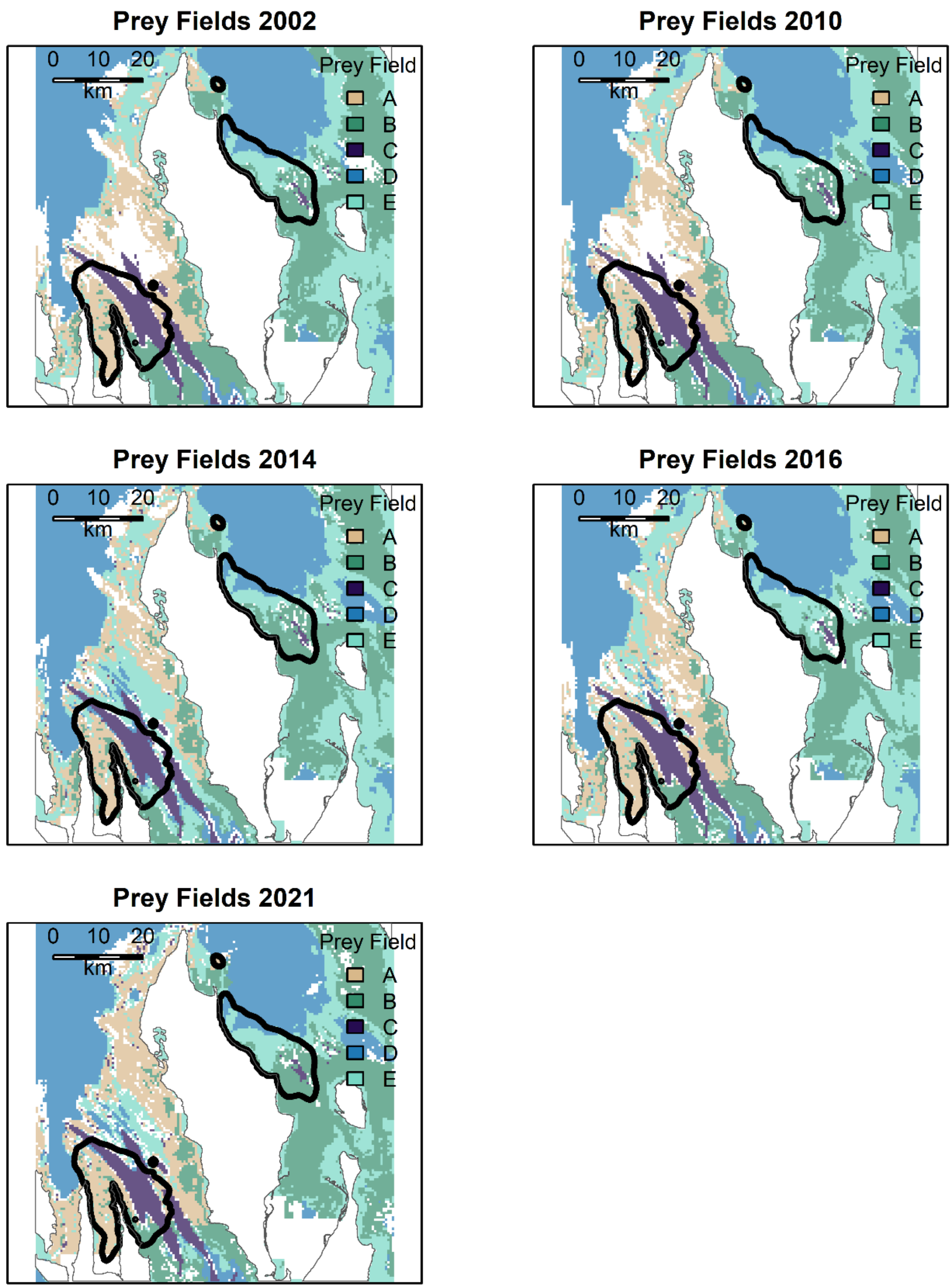


**Fig. S1.**
Time series of projected prey fields based on fish community distributions and varying seagrass coverage in Shark Bay, Western Australia. Projected prey field assignments are displayed only where assignment probability is $\geq 0.4$, lower values are shown as whitespace. Core study areas, defined by the 90% Kernel Density Estimate (KDE90), are outlined in black.

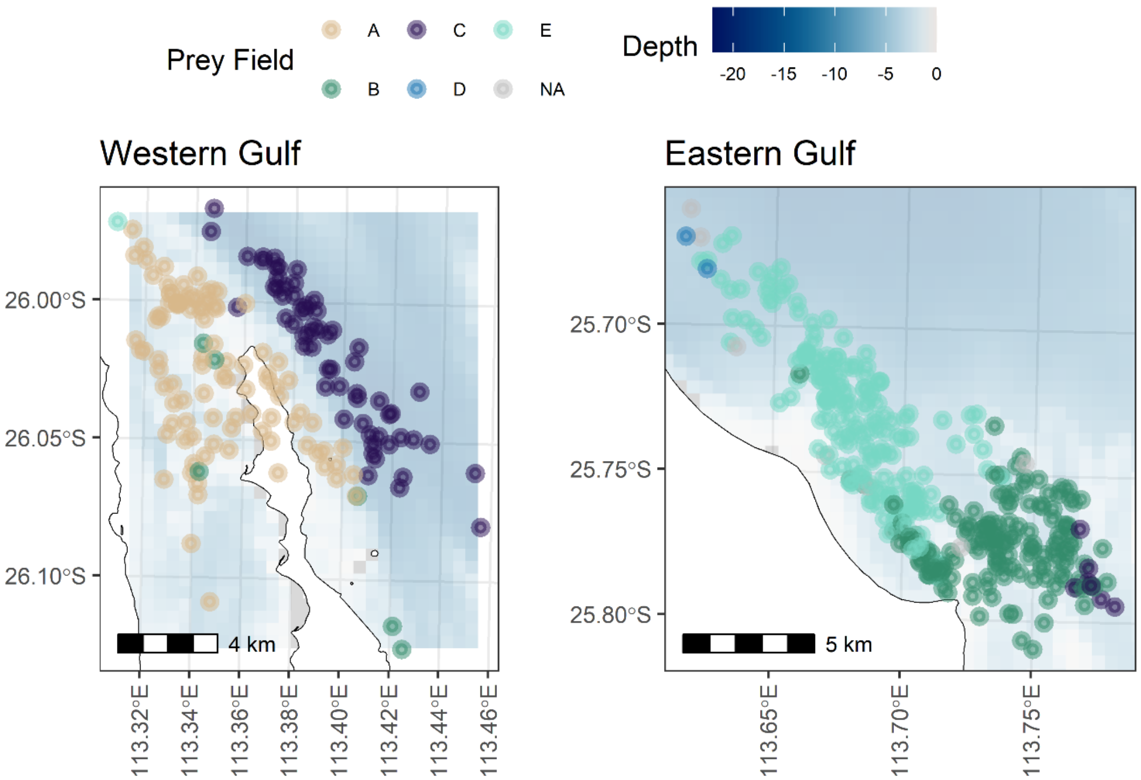


**Fig. S2.**

Overview of individual Indo-Pacific bottlenose dolphins (*Tursiops aduncus*) used for calculating the individual specialisation index ($S_I$) in Shark Bay, Western Australia. Landmasses are shown in white, while water bodies are shaded according to depth. Each point represents a dolphin, positioned at the centroid of all recorded foraging surveys for that individual, colour-coded according to the predominant foraging prey field of each dolphin. Points positioned on land are centroids of individuals recorded foraging in waters to the east and west of the land. Seven eastern gulf individuals could not be assigned a predominant prey field because they showed ties to two fields and are indicated with grey circles.

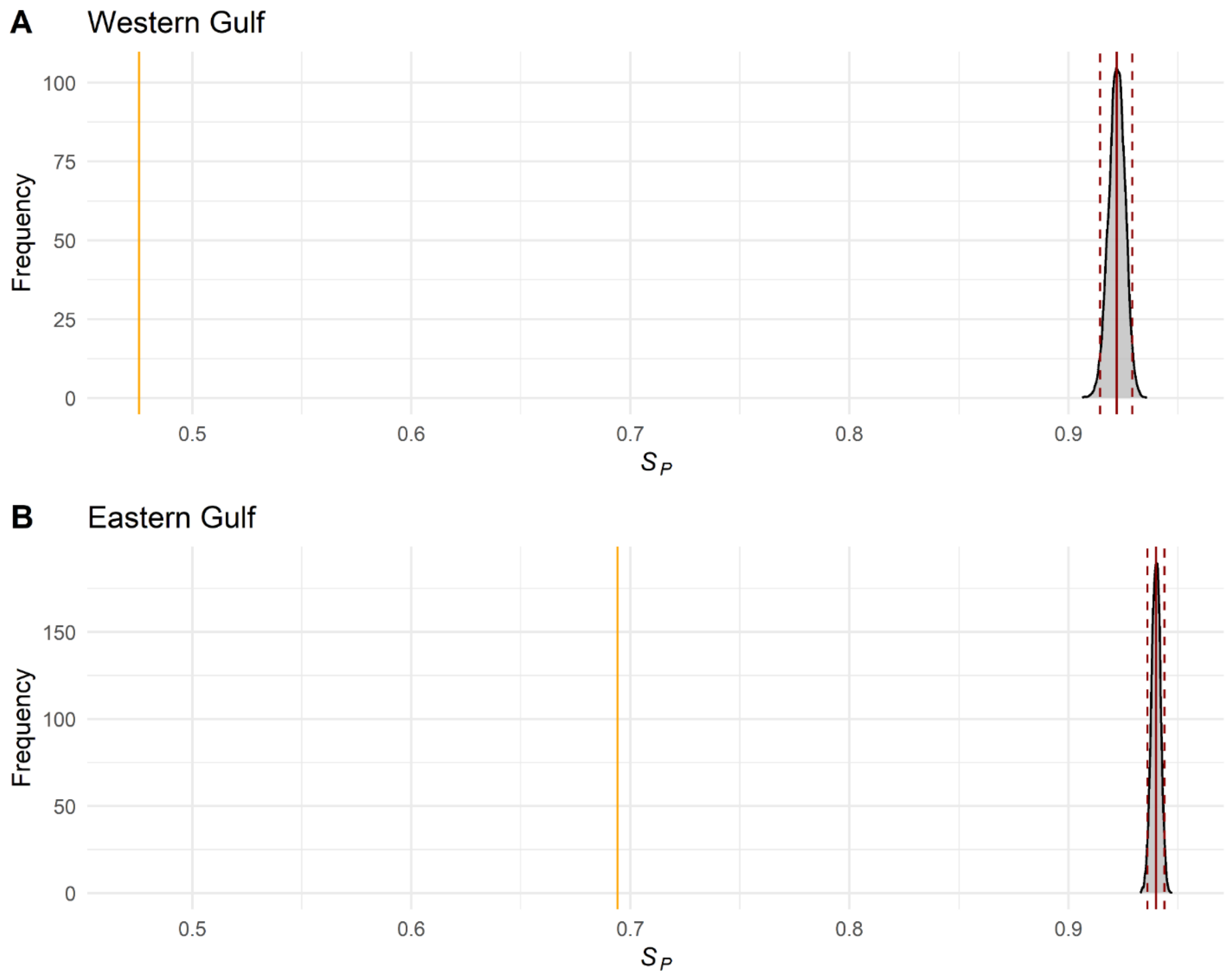


**Fig. S3.**

Distribution of random population level specialisation index ($S_P$) using Monte Carlo resampling with 9,999 permutations (grey). The mean and 95% confidence intervals are represented by solid and dashed red lines, respectively. The observed $S_P$ for subpopulation of Indo-Pacific bottlenose dolphins (*Tursiops aduncus*) in the western (**A**) and eastern gulf (**B**) of Shark Bay, Western Australia is shown as a solid orange line.

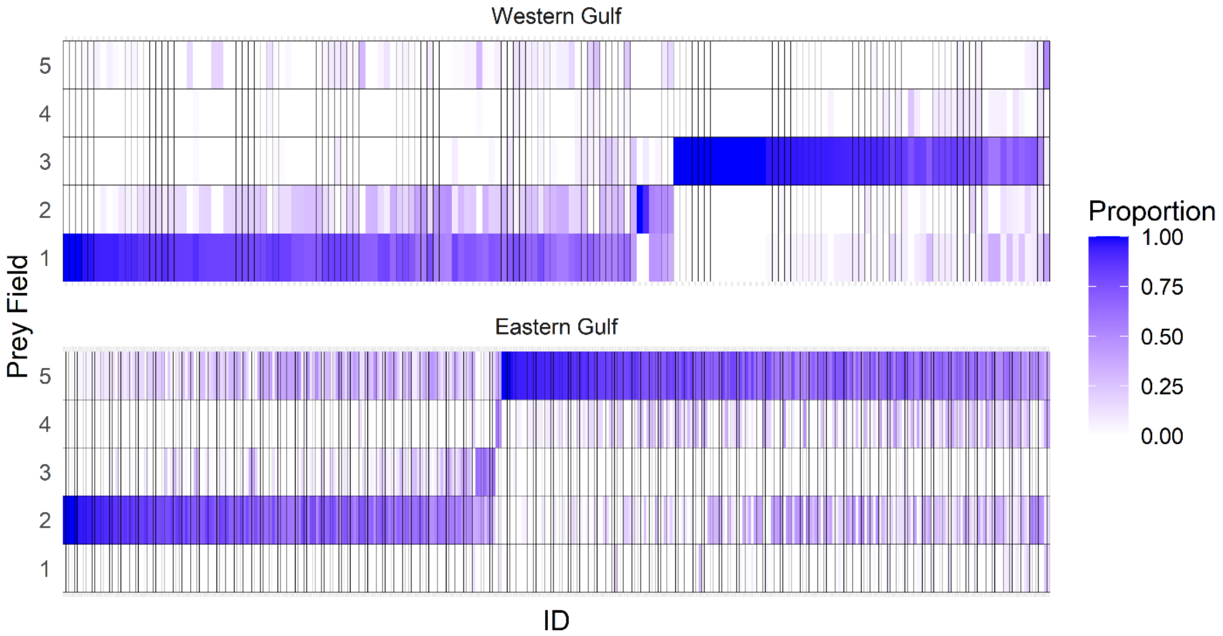


**Fig. S4.**

Foraging proportions of individual Indo-Pacific bottlenose dolphins (*Tursiops aduncus*) across different prey fields in the western and eastern gulfs of Shark Bay, Western Australia. Each row represents a prey field, while each column corresponds to an individual. The colour gradient indicates the observed proportion of foraging within each prey field, with darker shades representing higher proportions. Individuals are ordered first by their predominant foraging prey field and then by their strength of specialisation index ($S_I$). Prey field key: 1 = A, 2 = B, 3 = C, 4 = D, 5 = E

### Section S1

Comparison of individual prey field specialisation indices $S_I$ and age

We calculated the weighted mean age for each individual Indo-Pacific bottlenose dolphin (*Tursiops aduncus*) across all its foraging surveys to investigate whether $S_I$ was influenced by age. We used the age at the time of each foraging survey and calculated the mean across all surveys, such that individuals observed more frequently at younger ages received correspondingly lower mean ages. Our data included a weighted mean age range of 4.5–42.5 years (*mean*: 19.8 years), covering 361 individuals with available age estimates (western gulf: 27 females, 14 males; eastern gulf: 157 females, 164 males).

**Table S1.**

Bayesian generalized linear mixed model (GLMM) results for the effect of sex, subpopulation, and mean age on $S_I$: $R^2_c$= 0.14 (95% *CI* 0.09–0.20); $\Delta ELPD \pm SE$ = −21.7 ± 7.5 relative to null model.

| **Predictor** | ***β*** | **95% *CI*** |
|---|---|---|
| Subpopulation (western gulf vs. eastern gulf) | −0.29 | −0.39 – −0.19 |
| Sex (male vs. female) | 0.05 | −0.01 – 0.10 |
| Mean age | 0.00 | 0.00 – 0.00 |
| Log(survey count) | 0.04 | 0.00 – 0.08 |

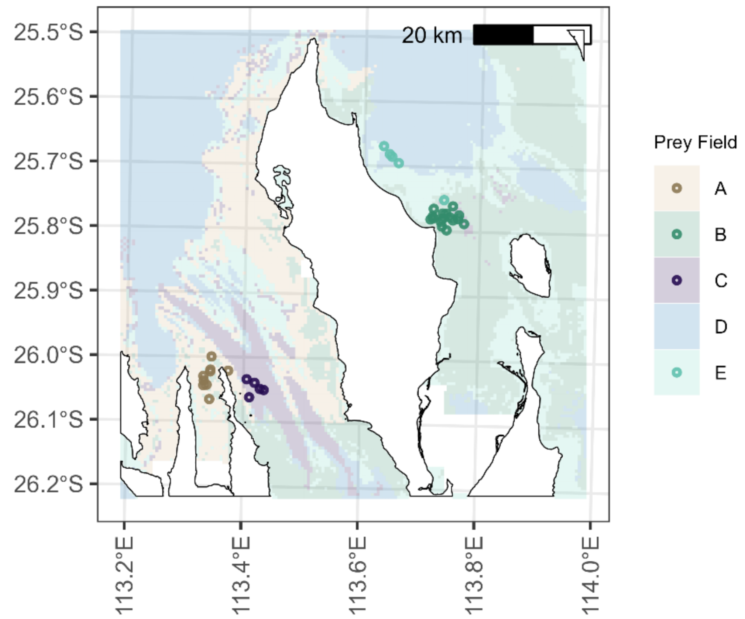


**Fig. S5.**

Spatial distribution of Indo-Pacific bottlenose dolphin (*Tursiops aduncus*) tissue samples collected from the western and eastern gulf of Shark Bay, Western Australia (points). Landmasses are shown in white, while water bodies are shaded according to their assigned prey fields according to the 2021 reference period. Points represent the centroid of all recorded foraging surveys for each sampled individual and are colour-coded according to their predominant foraging prey field.

**Section S2**

Yearly effects on stable isotope values

To rule out sampling year as a confounding effect, we modelled $\delta^{15}N$ and $\delta^{13}C$ as a function of time using all available stable isotope data ($N$ = 52), without restricting to individuals for whom a specialisation index could be calculated. For each isotope, we fitted a Bayesian GAM with a penalised spline on sampling year ($k$ = 7) to capture potential non-linear temporal trends, with Gaussian response distributions and weakly regularising priors (fixed effects: $\mu$ = 0, $\sigma$ = 5; intercept: $\mu$ = 0, $\sigma$ = 10; residual SD: Student's t, $\nu$ = 3, μ = 0, $\sigma$ = 10). Models were fitted using MCMC sampling with four chains run for 4,000 iterations for $\delta^{15}N$, and eight chains run for 10,000 iterations for $\delta^{13}C$ (including 2,000 warm-up iterations), the latter required to achieve adequate chain mixing. Model performance was evaluated as described in the main text. Neither temporal model improved predictive accuracy over the null model, and smooth terms ($\beta$) for year were not credibly different from zero in either isotope (Table S2), consistent with no meaningful interannual trend in $\delta^{15}N$ or $\delta^{13}C$.

**Table S2.**

Bayesian generalized additive model (GAM) results for temporal effects on $\delta^{15}N$ and $\delta^{13}C$ in skin samples of Indo-Pacific bottlenose dolphins (*Tursiops aduncus*) in Shark Bay, Western Australia.

| **Isotope** | ***ΔELPD*** | ***SE*** | ***β*** | **95% *CI*** | $R^2_c$ | **95% *CI*** |
|---|---|---|---|---|---|---|
| $\delta^{15}N$ | −0.1 | 2.0 | −1.59 | −5.90 – 1.79 | 0.10 | 0.005 – 0.27 |
| $\delta^{13}C$ | −1.0 | 1.1 | −0.43 | −5.84 – 3.31 | 0.05 | 0.002 – 0.17 |

### Section S3

Effect of prey field specialisation on $\delta^{13}C$ ratios

After ruling out interannual trends in stable isotope values (Section S2), we fitted a Bayesian GLM assessing the effect of each individual's predominant foraging prey field on $\delta^{13}C$ ratios, using the same modelling framework described in the main text. In contrast to $\delta^{15}N$, the model including predominant prey field, sex, and subpopulation did not outperform the null model, and no predictor showed a credible effect (Table S3; Fig. S6).

**Table S3.**

Bayesian generalized linear model (GLM) results for the effect of predominant prey field, sex, and subpopulation on $\delta^{13}C$: $\Delta ELPD \pm SE = -2.6 \pm 2.2$ relative to null model; $R^2_c = 0.17$ (95% *CI* 0.03–0.34) of Indo-Pacific bottlenose dolphins (*Tursiops aduncus*) in Shark Bay, Western Australia.

| **Predictor** | ***β*** | **95% *CI*** |
|---|---|---|
| Prey field B vs. A | −0.15 | −5.98 – 5.50 |
| Prey field C vs. A | −0.38 | −1.70 – 0.92 |
| Prey field E vs. A | −0.36 | −6.10 – 5.36 |
| Sex (male vs. female) | 0.43 | −0.33 – 1.25 |
| Subpopulation (WG vs. EG) | 0.50 | −5.14 – 6.12 |

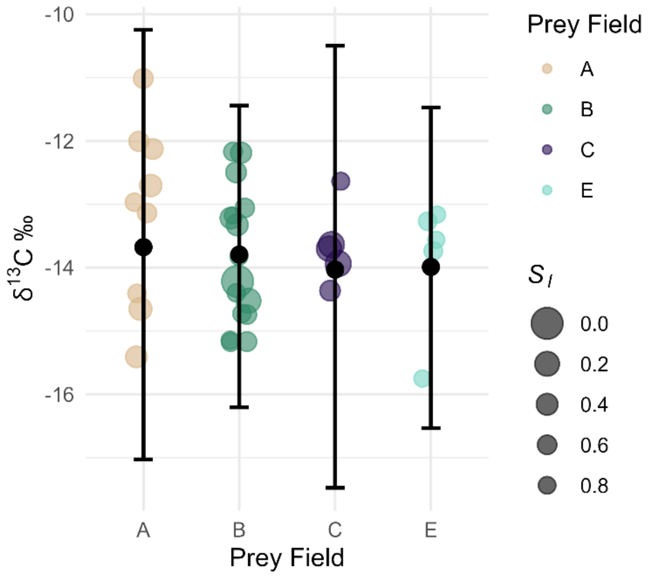


**Fig. S6.**

Relationship between predominant foraging prey field and $\delta^{13}C$ ratios in skin samples of Indo-Pacific bottlenose dolphins (*Tursiops aduncus*) in Shark Bay, Western Australia. Coloured points represent observed $\delta^{13}C$ ratios of individuals and their predominant foraging prey field. The size of the point indicates the strength of specialisation $S_I$. Black points and 95% credible interval whiskers represent model-predicted effects from the Bayesian generalized linear model (GLM) for $\delta^{13}C$ ratios fit with predominant foraging prey field as a predictor.

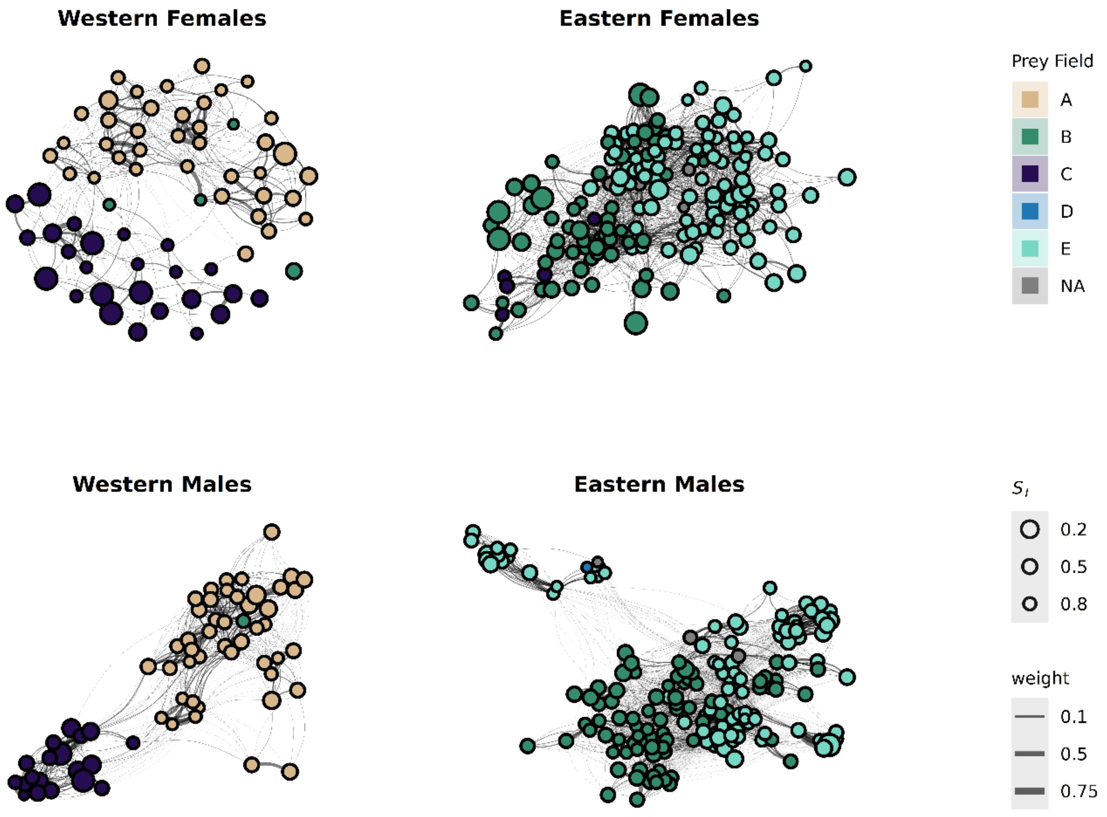


**Fig. S7.**

Social network plots for females and male Indo-Pacific bottlenose dolphins (*Tursiops aduncus*) from the western and eastern gulf subpopulations in Shark Bay, Western Australia. Individuals are represented as network nodes (points), which are connected by edges (lines) indicating social associations. Nodes are coloured according to the individual's specialisation in prey field and are sized based on the individual's degree of specialisation $S_I$. Edge widths represent dyadic social relationship indices *SRI*s, reflecting the strength of associations between individuals in non-foraging surveys.

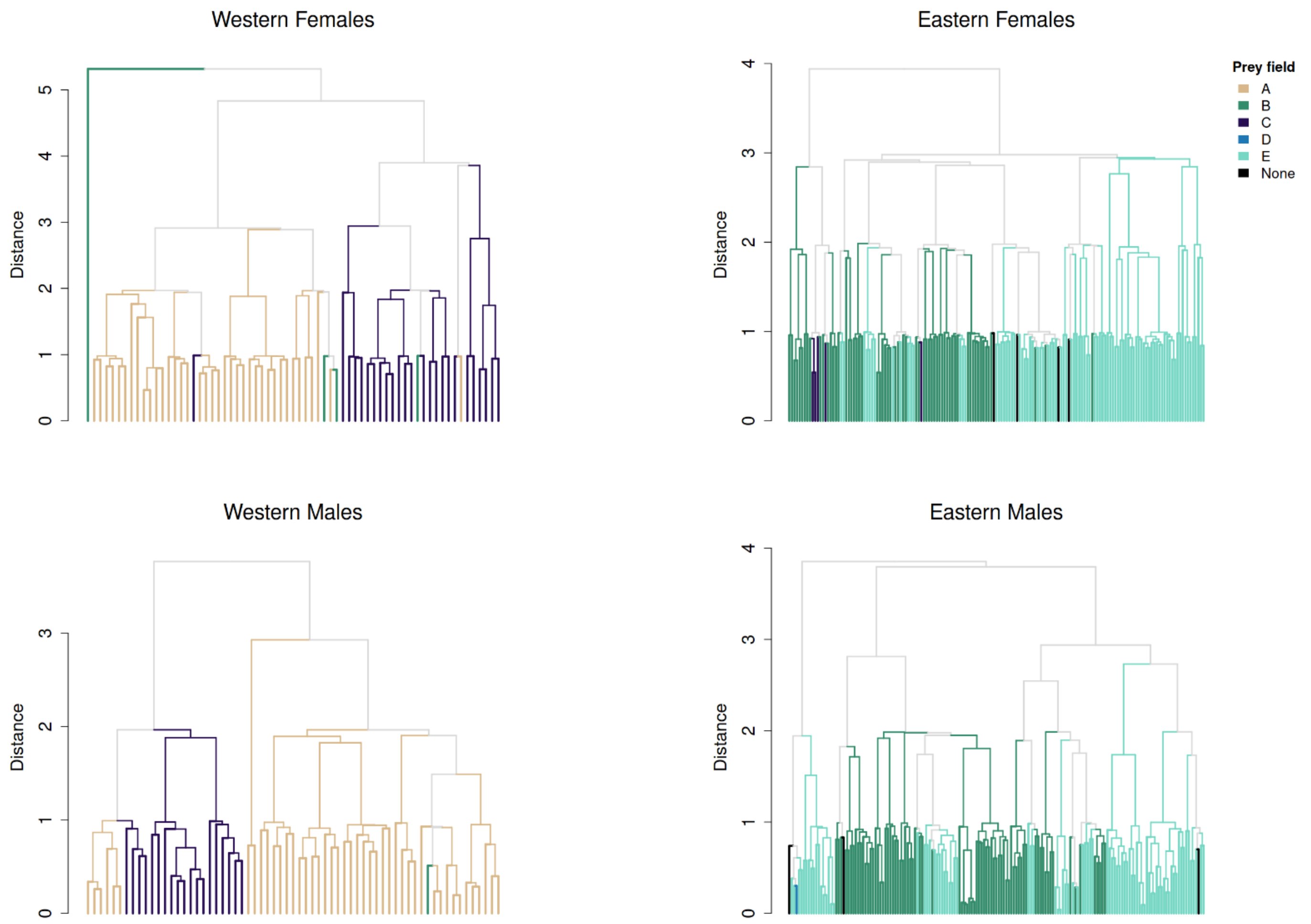


**Fig. S8.**

Hierarchical clustering of social networks by prey field specialisation for female and male Indo-Pacific bottlenose dolphins (*Tursiops aduncus*) from the western and eastern gulf subpopulations in Shark Bay, Western Australia. Dendrograms derived from complete-linkage hierarchical clustering of pairwise social distances (1 − Simple Ratio Index) for western gulf females, eastern gulf females, western gulf males, and eastern gulf males. Each leaf represents one individual, coloured according to their predominant foraging prey field (A–E; grey indicates clades with mixed prey field composition; black indicates individuals with no assigned prey field). The y-axis reflects social distance, such that individuals merging at lower values associate more strongly.

**Table S4.**

Summary of the Bayesian hurdle Gamma models linking foraging specialisation to betweenness centrality in non-foraging social networks of male and female Indo-Pacific bottlenose dolphins (*Tursiops aduncus*) in Shark Bay, Western Australia, in two subpopulations in the western and eastern gulf of the bay. Betweenness was modelled separately for each of the four sex–subpopulation networks because its distributional shape differed across networks in ways not addressable by standardisation alone. Each hurdle Gamma model includes $S_I$ and log-transformed non-foraging sighting frequency as predictors, with the gamma component (log link) modelling the magnitude of betweenness among bridge individuals and the hurdle component (logit link) modelling the probability of zero betweenness. Each model was compared via leave-one-out cross-validation against a null model including only log-transformed sighting frequency. $\beta$ values are posterior means with 95% credible intervals. Shape parameter estimates (gamma family): eastern gulf females = 0.83 (0.68 – 1.01); eastern gulf males = 0.62 (0.50 – 0.76); western gulf females = 1.08 (0.76 – 1.46); western gulf males = 0.78 (0.53 – 1.06). EG = eastern gulf, WG = western gulf. $N_{ind}$ = number of individuals in the network.

| **Network** | $N_{ind}$ | **Component** | **Parameter** | $\beta$ **(95% *CI*)** | $\Delta$***ELPD*** **±** ***SE*** | $R^2_c$ **(95% *CI*)** |
|---|---|---|---|---|---|---|
| **EG females** | 161 | **Gamma** | Intercept | 3.24 (2.22 – 4.29) | −0.1 ± 1.2 | 0.182 (0.062 – 0.327) |
| | | | $S_I$ | −0.24 (−0.91 – 0.42) | | |
| | | | log(surveys) | 0.61 (0.36 – 0.87) | | |
| | | **Hurdle** | Intercept | −0.20 (−2.33 – 1.99) | | |
| | | | $S_I$ | −0.78 (−2.05 – 0.49) | | |
| | | | log(surveys) | −0.33 (−0.90 – 0.20) | | |
| **EG males** | 169 | **Gamma** | Intercept | 4.74 (3.29 – 6.22) | 0.0 ± 1.2 | 0.068 (0.006 – 0.185) |
| | | | $S_I$ | 0.50 (−0.55 – 1.57) | | |
| | | | log(surveys) | 0.25 (−0.06 – 0.55) | | |
| | | **Hurdle** | Intercept | 0.94 (−0.95 – 2.86) | | |
| | | | $S_I$ | −0.76 (−2.03 – 0.52) | | |
| | | | log(surveys) | −0.38 (−0.83 – 0.05) | | |
| **WG females** | 67 | **Gamma** | Intercept | 2.37 (0.88 – 3.90) | −0.3 ± 0.7 | 0.121 (0.013 – 0.307) |
| | | | $S_I$ | 0.43 (−0.54 – 1.37) | | |
| | | | log(surveys) | 0.58 (0.11 – 1.06) | | |
| | | **Hurdle** | Intercept | −1.50 (−5.07 – 1.91) | | |
| | | | $S_I$ | 0.03 (−1.56 – 1.62) | | |
| | | | log(surveys) | −0.10 (−1.16 – 0.93) | | |
| **WG males** | 65 | **Gamma** | Intercept | 3.49 (1.23 – 5.88) | 0.0 ± 0.6 | 0.072 (0.004 – 0.215) |
| | | | $S_I$ | −0.63 (−1.81 – 0.55) | | |
| | | | log(surveys) | 0.47 (−0.19 – 1.10) | | |
| | | **Hurdle** | Intercept | 1.93 (−1.61 – 5.43) | | |
| | | | $S_I$ | 0.13 (−1.44 – 1.69) | | |
| | | | log(surveys) | −0.88 (−1.89 – 0.12) | | |

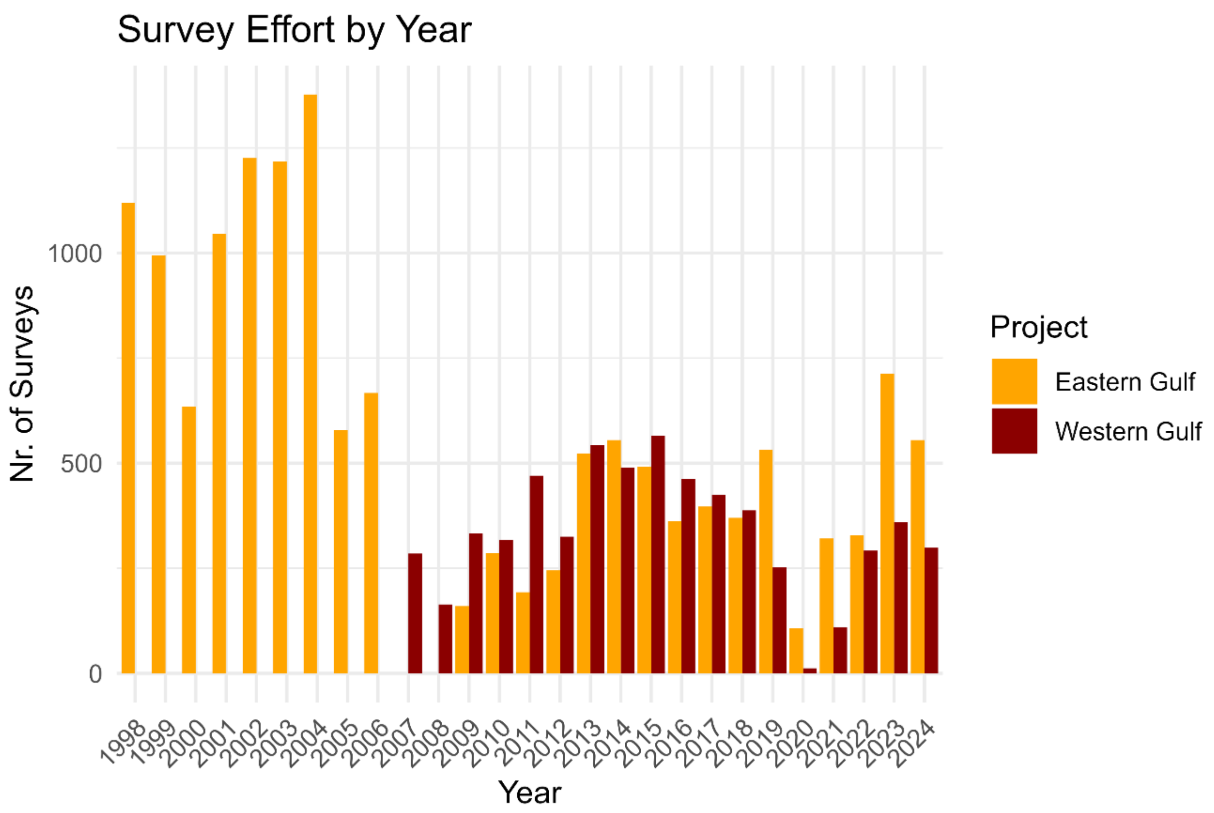


**Fig. S9.**
Total number of foraging surveys inside the core study sites (KDE90) per year and study site in Shark Bay, Western Australia, for Indo-Pacific bottlenose dolphins (*Tursiops aduncus*) in the eastern and western gulf.

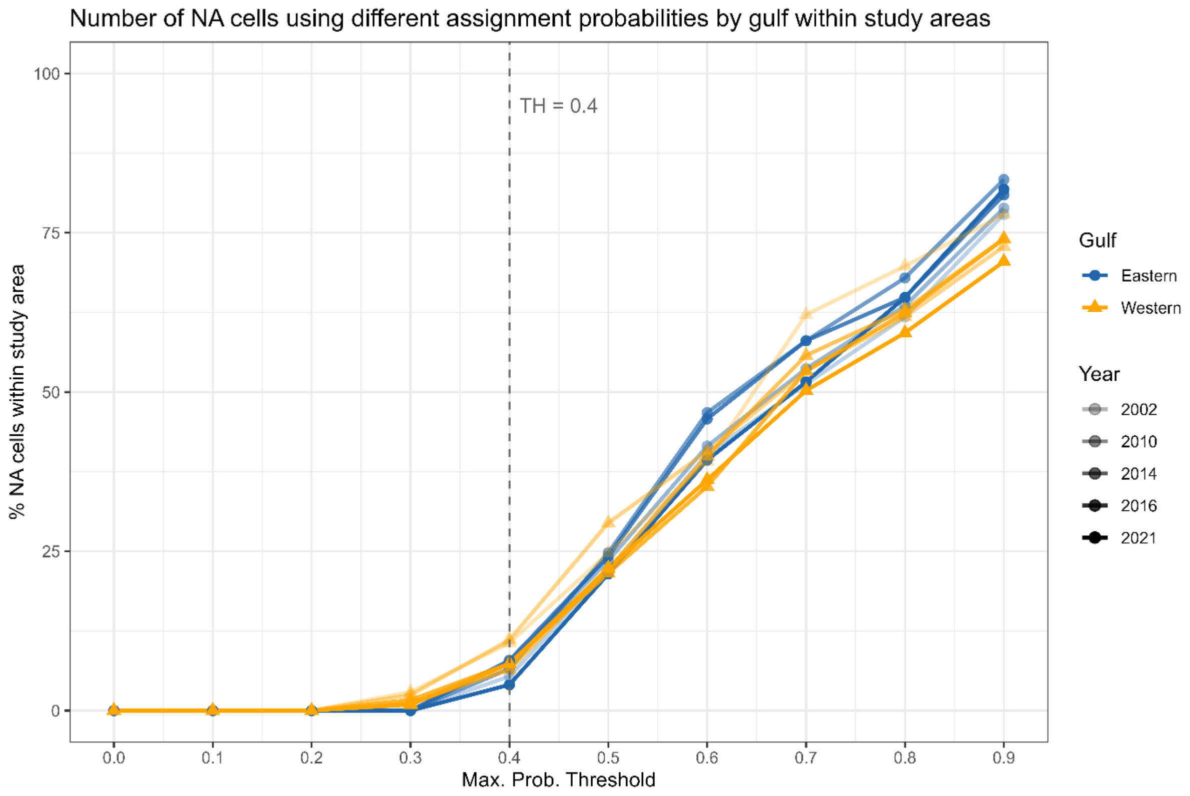


**Fig. S10.**

Sensitivity analysis of prey field assignment threshold for Indo-Pacific bottlenose dolphins (*Tursiops aduncus*) in Shark Bay, Western Australia. Percentage of unassigned grid cells (NA cells) within the KDE90 study areas of the eastern (blue) and western (orange) gulf as a function of maximum assignment probability threshold, shown separately for each of the five reference years (2002, 2010, 2014, 2016, 2021). The selected threshold of 0.4 (dashed vertical line) retained the majority of grid cells in both study sites across all reference years while excluding cells with low assignment confidence.

**Section S4**

Calculation of prey field specialisation indices

To calculate an index of specialisation (*S*) of foraging in a prey field, we applied Roughgarden's (1972) framework (*47*), which partitions the total niche width of a population (*TNW*) into within-individual (*WIC*) and between-individual (*BIC*) components.

According to (*80*), the degree of individual specialisation can be expressed as the proportion of *TNW* accounted for by within-individual variation:

$$S = \frac{WIC}{TNW} \tag{1}$$

where lower $S$ values indicate greater specialisation, and higher $S$ values suggest a more generalised foraging strategy.

To quantify *WIC* and *TNW*, we used the Shannon-Weaver index (*81*) with either the proportion of each individual’s surveys in each prey field, or the population’s surveys in each prey field, respectively (*80*).

More specifically, each individual’s within-individual component ($WIC_I$) was defined as:

$$WIC_I = \left(-\sum_j p_{ij} \ln p_{ij}\right) \tag{2}$$

The population-level $WIC_p$ was calculated as the weighted mean of all individual $WIC_I$ values:

$$WIC_p = \sum_i p_{i.} \left(-\sum_j p_{ij} \ln p_{ij}\right) \tag{3}$$

where

$$p_{ij} = \frac{n_{ij}}{\sum_j n_{ij}} \tag{4}$$

represents the individual’s proportion of foraging surveys in each prey field, and

$$p_{i.} = \frac{\sum_j n_{ij}}{\sum_i \sum_j n_{ij}} \tag{5}$$

was the proportion of an individual’s total foraging surveys relative to the entire dataset.

The *TNW* was calculated as:

$$TNW = -\sum_i q_j \ln q_j \tag{6}$$

Where

$$q_j = \frac{\sum_i n_{ij}}{\sum_i \sum_j n_{ij}} \qquad (7)$$

denoted the overall proportion of foraging surveys in each prey field.
Finally, each individual specialisation index, $S_I$ was calculated as according to eq. (1) with $S_I = WIC_I/TNW$, while the population-wide specialisation index was expressed as: $S_p = WIC_p/TNW$.

**Section S5**

Dyadic prey field overlap

As a measure of pairwise prey field overlap we used the Proportional Similarity Index (*$PS_I$*) (*80*), which measures the mean pairwise overlap between two individuals *i* and *k*:

$$PS_I = 1 - 0.5 \sum_j \left| p_{ij} - p_{kj} \right| \tag{8}$$

where $p_{ij}$ and $p_{kj}$ are the individuals i or k's observed proportion of foraging surveys in each prey field:

$$p_{ij} = \frac{n_{ij}}{\sum_j n_{ij}}; \; p_{kj} = \frac{n_{kj}}{\sum_j n_{kj}} \tag{9}$$

An $PS_I = 1$ indicates that the two individuals always forage in the same prey field or prey field composition, and $PS_I = 0$ indicates that the two individuals always forage in different prey fields.

**Section S6**

Simple Ratio Index

As a measure of social affiliation strength between two individuals, we used the Simple Ratio Index (*SRI*) (*86, 87*):

$$SRI = \frac{X}{X+Y_a+Y_b} \tag{10}$$

Where

$X$: The number of times the two individuals (dyad) were observed together.
$Y_a$: The number of times the first individual was observed without the second.
$Y_b$: The number of times the second individual was observed without the first.

An $SRI = 1$ indicates that the two individuals were always observed together and never apart, and $SRI = 0$ indicates that the two individuals were never observed together.
